# Mitosis without DNA replication in mammalian somatic cells

**DOI:** 10.1101/2020.07.08.193607

**Authors:** Olivier Ganier, Malik Lutzmann, Julien Cau, Isabelle Peiffer, Céline Lemmers, Sihem Zitouni, Charles Theillet, Marcel Méchali

## Abstract

DNA replication initiates with pre-replication complex (pre-RC) formation at replication origins in G1 (replication origin licensing), followed by activation of a pre-RC subset in the S phase. It has been suggested that a checkpoint prevents S phase entry when too few origins are licensed. Yet, we found that in normal cells, complete DNA synthesis inhibition by overexpression of a non-degradable geminin variant, or by CDT1 silencing prevents DNA replication without inducing any checkpoint. Cells continue cycling and enter mitosis, despite the absence of replicated DNA. Most of these unlicensed cells exit mitosis without dividing and enter senescence; however, about 25% of them successfully divide without previous DNA replication, producing daughter cells with half the normal diploid complement of chromosomes (1C). This suggests a potentially attractive strategy to derive haploid cells from any somatic cell type and unveil undescribed aspects of the coordination between DNA replication and cell division in mammals.

## INTRODUCTION

The tight coordination between DNA replication and cell division is regulated by several checkpoints to prevent premature mitotic entry before complete genome replication or in the case of persisting DNA damage^1,2,3,4,5^. Before DNA synthesis initiation, the spatial organization of DNA replication and the specification of replication origins are already set in the G1 phase of the cell cycle^6,7^. The assembly of the pre-replication complex (pre-RC) on DNA replication origins occurs in G1 (i.e. DNA replication origin licensing). The Origin Recognition Complex (ORC) binds to replication origins, followed by CDC6 and CDT1 recruitment that enables the loading of the replicative MCM2-7 helicase (reviewed in ^8^). Upon S phase entry, cyclin-dependent kinases (CDKs) activate replicative complexes, while inhibiting further licensing. This ensures that only one single round of DNA replication occurs per cell cycle. CDT1 contributes to prevent DNA replication re-initiation during S phase by restricting licensing to G1 (reviewed in ^9,8^). Upon replication fork activation, chromatin-bound CDT1 is degraded, and CDT1 binding to chromatin is prevented by geminin.

In mammalian cells, there is no example of cell division without DNA replication, except during meiosis. Examples are limited to the division of *Saccharomyces cerevisiae* haploid cells that results in the generation of cells with less than 1 C value^10,11,12^. In *S. cerevisiae*, the absence of licensing checkpoint might be linked to the lack of p53 and geminin^13,14,15^. Cell division without DNA replication can also be observed during early development of *Xenopus laevis*^16,17,18^ of and *Drosophila* embryos^19^, and was explained by the lack of cell cycle checkpoints at this early stage^17^.

Inhibition of DNA replication licensing for several days triggers G1-like arrest in normal cells, and massive death in cancer cells^20,21,13^. This G1-like block relies on the activation of p53 and p21; however, the sensitivity of this checkpoint remains unclear, particularly whether a complete licensing defect would be sensed and transduced to downstream effectors to halt cell cycle progression. Here, we show that DNA synthesis and mitosis are uncoupled when replication licensing is inhibited.

## RESULTS

### ΔDboxGeminin expression affects cell proliferation of normal cells without inducing a checkpoint response

We generated different stable cell lines where expression of a geminin mutant lacking the Destruction box (ΔDboxGeminin) is induced by doxycycline (DOX) (Supplementary Figure 1A). As ΔDboxGeminin cannot be degraded, its expression is stabilized in the G1 phase (Supplementary Figure 1B), while endogenous geminin is degraded upon APC/C action^22^. Upon DOX addition, ΔDboxGeminin expression was induced in 82% of cells (Supplementary Figure 1C).

The long-term effect of ΔDboxGeminin expression differed in normal somatic cells and in cancer cells, as previously reported^23,24^. In the tested cancer cell lines (HCT116, T98G, HeLa and HEK 293 cells), ΔDboxGeminin expression led to massive cell death, as indicated by the cell number decrease and the appearance of a spread sub-G1 population (cytometry analysis) 72 hours after induction (+DOX) (Figure 1A and Supplementary Figure 1D-E). In untransformed cells, such as NIH 3T3 fibroblasts and adult tail tip fibroblasts (TTFs), ΔDboxGeminin expression did not induce cell death, but blocked cells in a G1-like phase, as suggested by the accumulation of cells with 2C DNA content (2C cells) (Figure 1A and Supplementary Figure 1D), confirming previous findings^23,24^.

**FIGURE 1:**
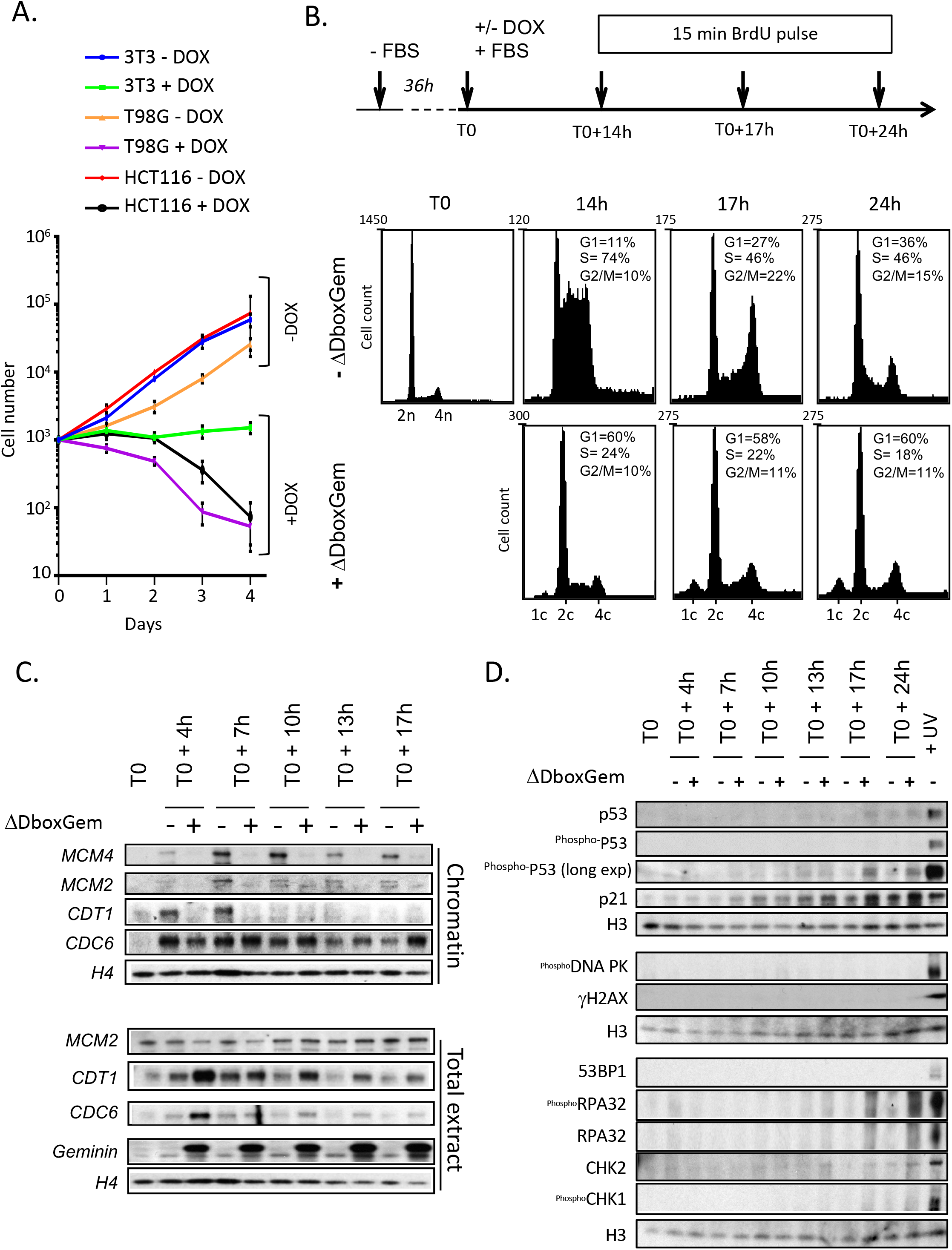
Induction of ΔDboxGeminin blocks DNA replication licensing without inducing the intra-S phase or DNA damage checkpoints in somatic cells. **A.** Δ*DboxGeminin induction inhibits cell proliferation in untransformed somatic cells and kills cancer cells*. Cell proliferation analysis in 3T3 fibroblasts (3T3), HCT116 (p53^+/+^) colorectal cancer cells and p53-deficient T98G brain glioblastoma cells before (−) and after (+) induction of ΔDboxGeminin expression by incubation with doxycycline (DOX). **B.** *ΔDboxGeminin inhibits DNA replication*. Upper panel. Scheme of ΔDboxGeminin induction in 3T3 fibroblasts. Cells were synchronized in G0 (T0) by serum starvation (-FBS) and released by FBS addition in the medium with or without (+/−) DOX to induce ΔDboxGeminin expression. Cells were then incubated with BrdU before harvesting. Lower panels. Fluorescence-activated cell sorting (FACS) analysis of 3T3 cells incubated or not with DOX (+/− ΔDboxGeminin) after BrdU addition for 15 minutes before collection at 14h, 17h and 24h after release. The x-axis shows the DNA content (propidium iodide staining) and the y-axis the cell number in each cell cycle phase, determined by BrdU incorporation and DNA content. **C.** *ΔDboxGeminin expression prevents DNA replication licensing by blocking CDT1 action.* Chromatin-bound (upper panels) and total proteins extracts (lower panels) were obtained from 3T3 cells (+/− DOX) at different time points after release from G0 and analysed by western blotting. CDT1 binding to chromatin was inhibited by ΔDboxGeminin expression, but not its accumulation in total protein extracts, confirming that CDT1 degradation is dependent on DNA replication and not on cell cycle progression. Binding of MCM2 and MCM4 to chromatin also was inhibited upon DOX addition, as expected in cells with a licensing reaction defect. Conversely, CDC6 binding to chromatin was not affected, because it occurs before and independently of CDT1. **D.** *Licensing inhibition by ΔDboxGeminin does not induce checkpoint activation during the first cell cycle.* Different checkpoint proteins were analysed by western blotting after licensing inhibition. As a positive control, 3T3 cells were exposed to UV to confirm that checkpoints are functional in these cells.

As we observed these phenotypes after several days of replication licensing block, we wanted to decipher the earliest mechanisms leading to this cell cycle arrest in normal cells. Therefore, we synchronized non-transformed somatic 3T3 cells and induced ΔDboxGeminin expression upon release from serum starvation (Figure 1B). ΔDboxGeminin induction prevented CDT1 binding to chromatin in G1, without any effect on CDT1 abundance (Figure 1C) and on chromatin binding of pre-RC components that act upstream of CDT1, such as CDC6. Conversely, it dramatically reduced the chromatin association of replication factors downstream of CDT1, such as MCM2 and MCM4, confirming that the licensing step was inhibited (Figure 1C). FACS analyses (Figure 1B) also confirmed that S phase progression was inhibited (24% vs 74% of BrdU^+^ cells in non-induced cells, 14 hours after release). The presence of replicating (BrdU^+^) cells upon DOX addition can be readily explained by the fraction of cells in which DOX did not induce ΔDboxGeminin expression, as confirmed by FACS analysis (Supplementary Figure 1C). At 17 hours post-release, 58% of ΔDboxGeminin-expressing cells were still in a G1-like state (2C DNA content and no BrdU incorporation) (Figure 1B).

As cell cycle arrest *via* p53 and p21 induction was observed several days after inhibition of DNA replication licensing^23,24,25^, we asked whether p53 and p21 induction could represent a later indirect consequence of cell cycle arrest. ΔDboxGeminin expression in G1 resulted in efficient inhibition of licensing and DNA replication, but we detected p53 phosphorylation and p21 upregulation (markers of cell cycle arrest) only at late time points (Figure 1D, see T0+17h), when control cells (no DOX) had already completed mitosis and entered the next cell cycle (Figure 1B, T0+17h). We concluded that induction of p53 and p21 was a late indirect event induced upon licensing inhibition in non-cancer cells.

Conversely, during the first cell cycle following ΔDboxGeminin induction, the intra-S phase checkpoint, the G2-M checkpoint (induction of CHK1 or CHK2) and the DNA damage checkpoint (phosphorylation of RPA32, H2AX and DNA-PK) were not induced in unlicensed cells (Figure 1D). Induction of p53 upon UV-induced DNA damage confirmed the integrity of the p53 pathway in these cells (Figure 1D).

We concluded that ΔDboxGeminin expression in mammalian somatic cells prevents origin licensing and inhibits DNA replication, without checkpoint induction during the first cell cycle.

### Mitotic progression of somatic cells in the absence of DNA replication licensing

Unexpectedly, ΔDboxGeminin-expressing cells continued cycling. Markers of cell cycle entry showed similar kinetics in ΔDboxGeminin-expressing and control cells (Figure 2A). Compared with T0 (release from G0), the CDK inhibitor p27 was progressively downregulated and pRB phosphorylation increased, indicating cell cycle progression. Likewise, cyclin A2, a marker of S-phase entry, was normally induced, as well as markers of the G2-M phases, such as cyclin B1, PLK1 and phosphorylation of histone H3 on serine 10 (^ser10^PH3) (Figure 2A). These results confirmed the absence of checkpoints to block cell cycle progression after origin licensing inhibition (Figure 1D).

**FIGURE 2:**
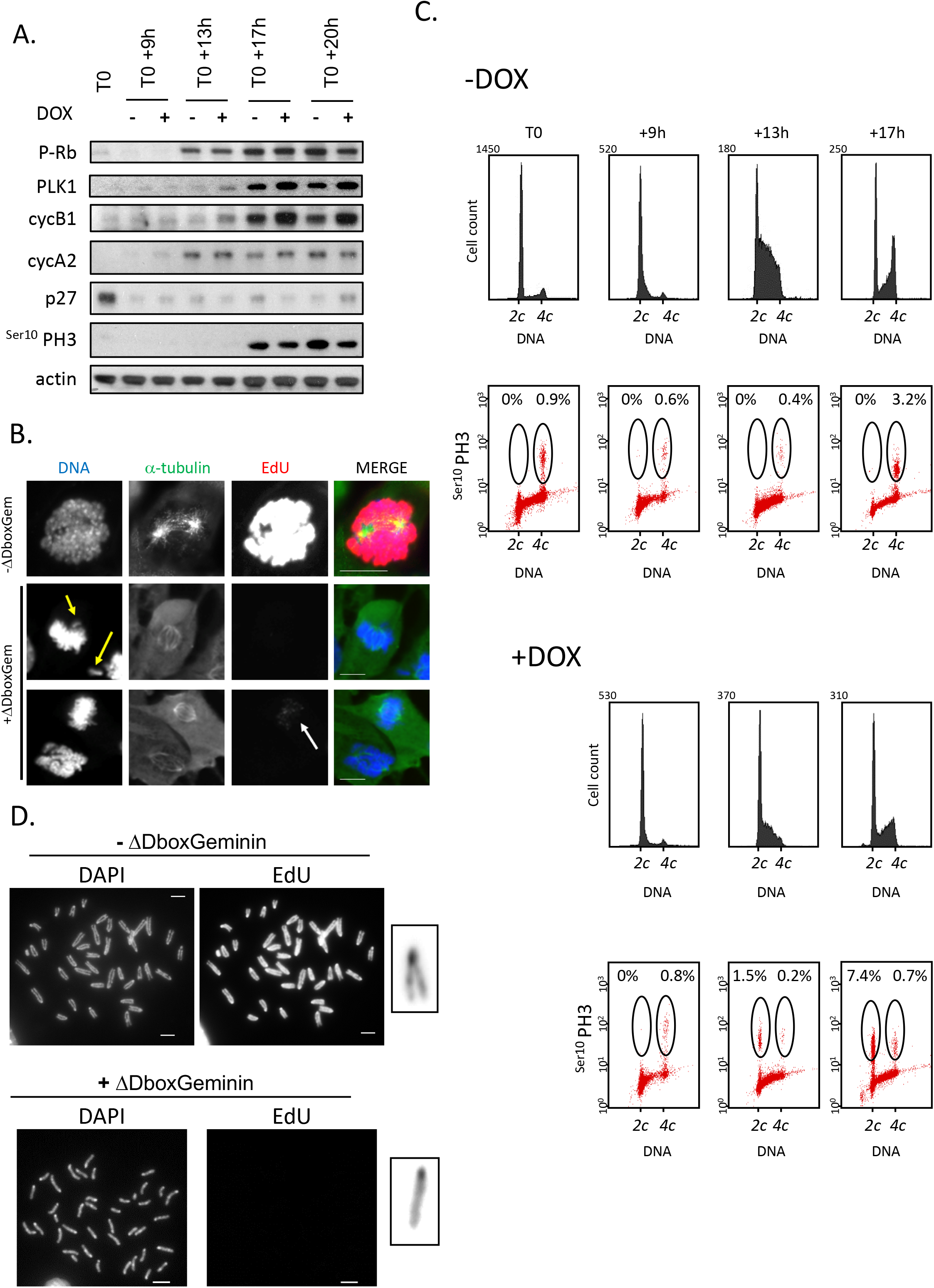
In somatic cells, expression of ΔDboxGeminin does not block cell cycle progression and induces mitotic entry of unlicensed cells. **A.** *ΔDboxGeminin expression does not prevent the expression of mitotic markers.* Total protein extracts from 3T3 cells were analysed by western blotting at different time points after release from quiescence (G0=T0) and induction (+) or not (−) of ΔDboxGeminin expression by DOX. **B.** *Immunofluorescence analysis of control (-ΔDboxGem) and unlicensed mitotic cells (+ΔDboxGem).* 3T3 cells were synchronized in G0 and then released in the presence of EdU (+/− DOX to induce ΔDboxGeminin expression). At T0+20h, cells were fixed to assess EdU incorporation and stained with antibodies against α-tubulin to visualize the mitotic spindle. Yellow arrows show lagging chromosomes in unlicensed (EdU negative) mitotic cells. Cells in which licensing was partially inhibited could also enter mitosis, as indicated by the incomplete EdU staining of mitotic chromosomes (white arrow). Scale bars = 10 μm. **C.** *ΔDboxGeminin expression leads to mitotic entry of cells with 2C DNA content*. ^ser10^PH3 signal in 3T3 cells that do not (-DOX, upper panels) and that express (+DOX, lower panels) ΔDboxGeminin and that have 2C (pre-replicative cells) and 4C (post-replicative cells) DNA content, assessed by flow cytometry at different times after release from quiescence. DNA content was evaluated after propidium iodide staining and the percentage of 2C and 4C cells is indicated for each condition. **D.** MEFs that express inducible ΔDboxGeminin were synchronized in G0 and released in the presence of EdU with (+) or without (−) DOX to induce or not ΔDboxGeminin expression. After 12 hours, cells were incubated with colcemid for 4 hours and EdU incorporation was assessed in metaphase spreads. Control metaphase spreads (−ΔDboxGeminin) displayed 40 chromosomes with two replicated chromatids attached by the kinetochore. Conversely, some metaphase spreads of ΔDboxGeminin-expressing cells (+ΔDboxGeminin; example in the left panels) were EdU-negative and had 40 chromosomes with single chromatids (see enlarged inset). Scale bars = 8 μm.

We then asked whether unlicensed cells could progress to mitosis. We synchronized 3T3 fibroblasts in G0/G1 and released them in the presence of EdU to monitor DNA replication (Supplementary Figure 2A). Strikingly, in ΔDboxGeminin-expressing cells, we observed unreplicated mitotic cells with normal chromatin compaction and bipolar spindles (Figure 2B and Supplementary Figure 2) and positive for the mitotic marker ^ser10^PH3, but without detectable EdU signal. We never observed this phenotype in control cells.

We concluded that in the absence of DNA replication licensing, the cell cycle can progress until mitosis.

### Mitotic cells with 2C DNA content are generated after licensing inhibition

To determine whether the mitotic cells observed in the absence of DNA replication harbour a 2C genome, we analysed their DNA content. As expected, control cells expressed ^ser10^PH3, a mitosis marker, only after DNA replication (4C cells), but never in pre-replicative 2C cells (Figure 2C). Conversely, in ΔDboxGeminin-expressing cells, we observed ^ser10^PH3 also in 2C cells, as expected for unreplicated mitotic cells (Figure 2C). We confirmed the mitotic status of these 2C cells using other mitotic markers (Supplementary Figure 3). We also observed mitotic entry of unlicensed cells when using primary somatic mammalian cells (mouse embryonic fibroblasts, MEFs, and adult TTFs) and in cancer cells (Supplementary Figure 4). Moreover, a three-color flow cytometry approach showed that in ΔDboxGeminin-expressing 2C cells, ^ser10^PH3 could be detected in EdU-negative cells (Supplementary Figure 5).

Of note, when ΔDboxGeminin expression was induced in cells already progressing through S phase, the first cell cycle was not affected, as expected for a G1-specific effect of ΔDboxGeminin on DNA replication licensing. Using this experimental setting, 2C mitotic cells were generated during the second cell cycle (Supplementary Figure 6). This observation shows that mitotic entry of unreplicated cells after licensing inhibition by ΔDboxGeminin was not restricted to the first cell cycle following release from quiescence ^25^.

To confirm that ΔDboxGeminin effect was through CDT1 inhibition^26^, we downregulated CDT1 by RNA interference (shCDT1). We detected a ^ser10^PH3-positive 2C cell population in shCDT1 cells and confirmed the mitotic entry of non-replicated shCDT1 cells by immunofluorescence (Supplementary Figure 7).

Finally, we asked whether mitotic chromosomes were formed in 2C mitotic cells. We prepared metaphase spreads of MEFs 10 hours after synchronization in G0/G1 and release in the presence of EdU. Control cells had 40 pairs of chromosomes with the two replicated (EdU-positive) sister chromatids attached by the kinetochore (Figure 2D). Conversely, ΔDboxGeminin-expressing cells were EdU-negative and had 40 single-chromatid chromosomes. This unequivocally demonstrated the uncoupling of DNA replication and mitosis after licensing inhibition in mammalian (normal and transformed) cells.

### A discrete population of unlicensed cells can successfully divide

We then asked whether unlicensed cells could complete cell division. We synchronized 3T3 cells in G0/G1, and then released them in the presence of DOX and EdU to monitor DNA synthesis (Figure 3A). We used time lapse microscopy to follow mitosis progression (Supplementary movies 1 and 2), and then fixed cells to check whether DNA had replicated before mitosis. The used cell lines also expressed GFP-tagged histone 2B (GFP-H2B), a chromatin marker.

**FIGURE 3:**
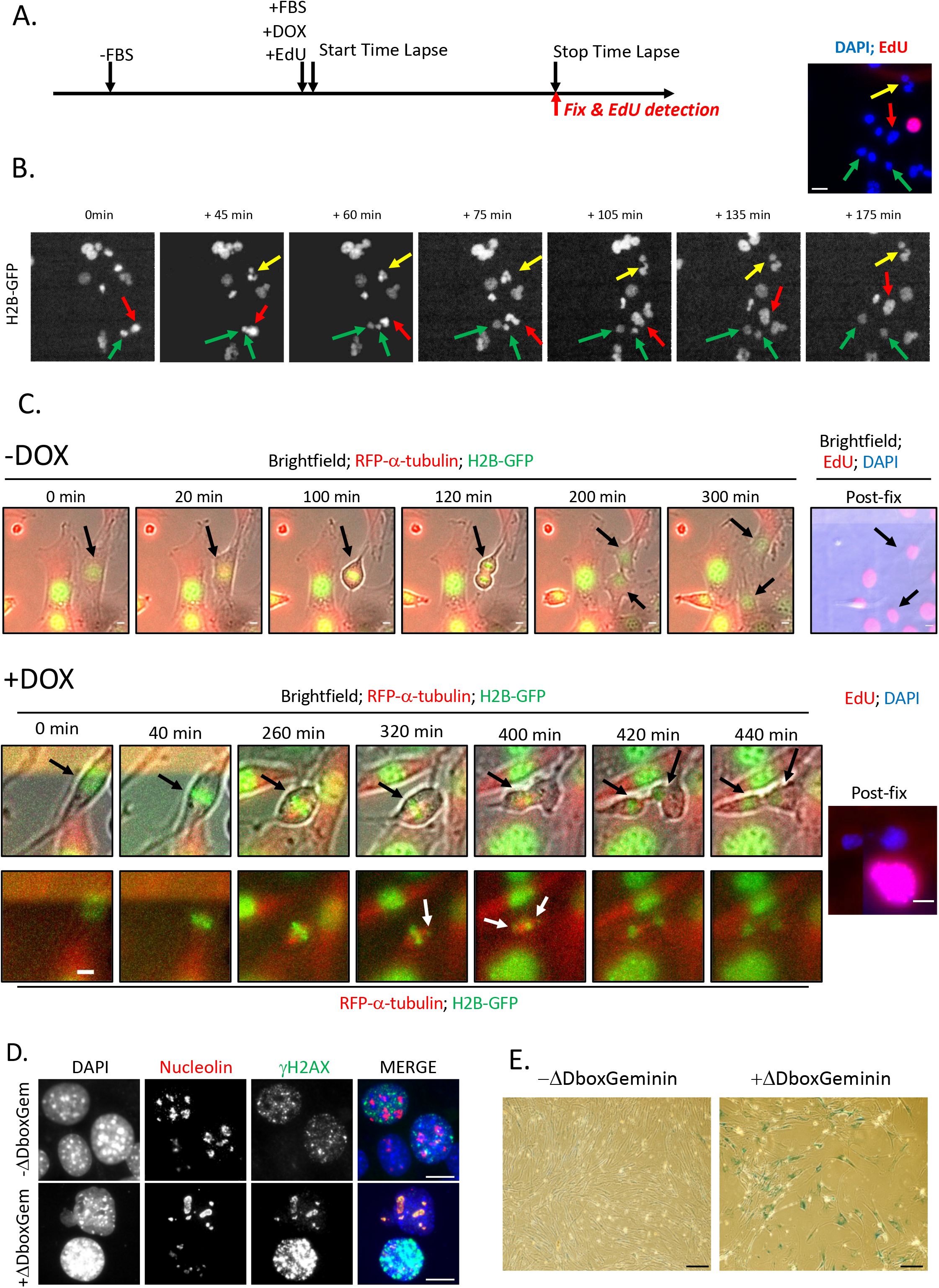
Unlicensed mitotic cells can generate two daughter cells despite the absence of DNA replication. **A.** Schematic representation of the strategy to investigate the fate of unlicensed (EdU-negative) mitotic cells. 3T3 cells that transiently express GFP-H2B were synchronized in G0 and released in the presence of EdU and DOX to induce ΔDboxGeminin expression. At the end of the time lapse microscopy recording, cells were fixed and processed for EdU detection. **B.** *Images of a time-lapse experiment showing the different fates of unlicensed mitotic cells*: i) completion of cell division (green arrows); ii) formation of a bi-nucleated cell (cytokinesis failure) (yellow arrows); and iii) formation of a mono-nucleated cell (mitotic slippage) (red arrows). Images were acquired every 15 minutes and the time stamps are indicated. Lower panels: fluorescence images; upper panel: corresponding cells fixed at the end of the time lapse and after EdU (red) and DAPI (DNA; blue) staining. Scale bars = 25 μm. **C.** *Unlicensed cells can divide without DNA replication.* Still images from the time-lapse recording of the experiments described in A and showing a control cell (upper panels, -DOX) and a ΔDboxGeminin-expressing cell (lower panels, +DOX) that express H2B-GFP (green) and RFP-α-tubulin (red) and are undergoing mitotic division. Images were acquired every 20 minutes and the time stamps are indicated. Black arrows show the mitotic cell and then its two daughter cells. The right panels show the EdU status of the two daughter cells. White arrows indicate the lagging chromosomes that can be transiently observed before cell division completion of the unlicensed mitotic cell. Scale bars = 10 μm. **D.** 24 hours after release from G0 and incubation with (+ΔDboxGem) or without (- ΔDboxGem) DOX, 3T3 cells were fixed and the expression of γH2AX and nucleolin (a marker of nucleoli) was assessed by immunofluorescence. Scale bars = 5 μm. **E.** 10 days after ΔDboxGeminin induction by incubation (+ΔDboxGeminin) or not (- ΔDboxGeminin) with DOX, mouse adult tail-tip fibroblasts were fixed and stained to monitor β-galactosidase activity, a marker of premature senescence. Scale bars = 20 μm.

Time lapse experiments revealed that 76% of EdU-negative mitotic cells did not divide and resulted in bi-nucleated cells (cytokinesis failure, Figure 3B, yellow arrow) or in cells with a single nucleus (mitotic slippage, Figure 3B, red arrow).

Overall, 24% of unlicensed cells that entered mitosis could successfully divide (EdU-negative, green arrows in Figure 3B, Supplementary movies) despite the occasional presence of lagging chromosomes (Figure 3C, white arrows, lower panel). Importantly, mitosis duration was longer in unlicensed cells than in control cells (Figure 3C), explaining the accumulation of mitotic cells observed by flow cytometry (Figure 2).

In summary, although the majority of unlicensed cells could not complete cell division, about 25% of them produced two viable daughter cells.

### Mitotic division failure in unlicensed mitotic cells leads to long-term irreversible G1 arrest with senescence features

We further investigated the fate of normal 3T3 cells that entered mitosis without previous DNA replication, but did not complete cell division. The first wave of mitotic cells started at 14h post-release with a peak at 17h (Supplementary Figure 8). The percentages of anaphase bridges and micronuclei after the beginning of mitotic entry were higher in ΔDboxGeminin-expressing than control cells (Supplementary Figure 8A-B). This confirmed the mitosis defects of unlicensed mitotic cells observed by time lapse microscopy (Figure 3B). Moreover, 48h after induction, ΔDboxGeminin-positive cells showed unusually large and weak foci of phosphorylated H2AX (γH2AX) that decorated nucleoli after the first mitotic wave, when p21 and p53 were induced (Figure 3D, Supplementary Figure 8C-D). These persistent nucleolar foci were evocative of those observed in old hematopoietic cells undergoing senescence ^27^. Indeed, cells with nucleolar γH2AX foci were always EdU-negative (Supplementary Figure 8C), suggesting that they had exited the cell cycle.

At day 14 after ΔDboxGeminin induction, most cells were still alive, but had entered premature senescence, as revealed by β-galactosidase staining (Figure 3E).

We concluded that unlicensed mitotic cells that did not divide were subsequently blocked in G1. This was associated with atypical nucleolar DNA damage, induction of the DNA damage checkpoint and senescence features that only appear after the first mitotic wave. Therefore, p53 and p21 were induced in unlicensed cells, as previously reported^23,25^; however, this effect was not primarily triggered by the lack of licensed origins, but by mitotic failure-induced DNA damage, as observed after abortive completion of normal mitosis (i.e. 4C DNA content, post-replicative cells)^28–31^.

### Division of unlicensed cells leads to a cell population with 1C DNA content

The fraction of unlicensed cells (24%) that divided produced two daughter cells, despite the complete absence of DNA replication. If the unlicensed genomes were equally distributed in the two daughter cells, these cells should have 1C DNA content. In agreement, flow cytometry analyses always highlighted a distinct population of ΔDboxGeminin-expressing cells with 1C DNA content that appeared after the first cell cycle (Figures 1A and 4A). We confirmed that they were daughter cells (i.e. the result of mitosis) because inhibition of mitotic entry by incubation with the CDK1 inhibitor RO-3306 totally prevented their appearance (Figure 4A). We could not detect cleaved caspase 3 (a marker of apoptosis), unlike in control UV-irradiated cells (Figure 4B). Moreover, after UV exposure, cell cycle progression of unlicensed cells was inhibited by the DNA damage checkpoint (as indicated by p53 phosphorylation; Figure 4C), preventing the appearance of 1C cells (Figure 4A). Noticeably, the sub-G1 population of ΔDboxGeminin-expressing cells (with 1C DNA content) showed a sharp profile, as opposed to the classic spread-out profile of dying sub-G1 cells after UV irradiation (Figure 4A).

**FIGURE 4:**
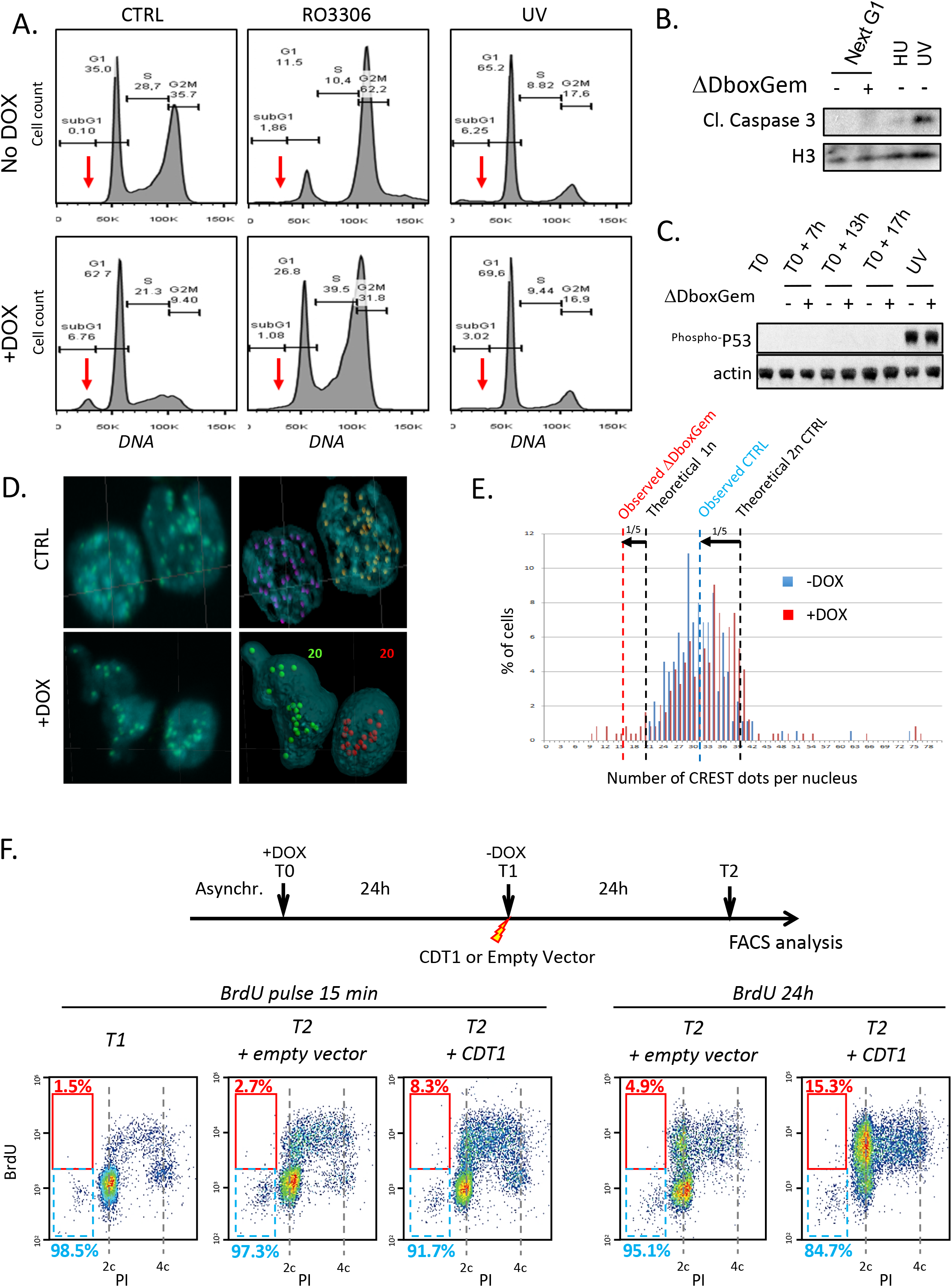
Completion of mitosis in unlicensed cells leads to a viable cell population with 1C DNA content that can re-enter the cell cycle. **A.** FACS profiles showing the DNA content (propidium iodide staining) of NIH 3T3 cells 20 hours after release from G0 in the presence (+) or not (−) of DOX alone (CTRL) or combined with RO-3306 (a CDK1 inhibitor that prevents mitotic entry), or after exposure to UV upon release to induce DNA damage. The sub-G1 population (red arrow) observed in ΔDboxGeminin-expressing cells (bottom left panel) showed a discrete and relatively sharp profile. RO-3306 prevented mitotic entry and the appearance of the 1C DNA cell population (middle bottom panel). UV treatment blocked cell cycle progression in control cells (no DOX) (top right panel) and induced apoptosis, as indicated by the spread sub-G1 population. UV exposure prevented the appearance of the 1C cell population in ΔDboxGeminin-expressing cells (bottom right panel), showing that the DNA damage checkpoint is intact in these cells and can block progression through the cell cycle of unlicensed cells (and therefore mitosis). **B.** Western blot analysis confirmed the presence of apoptotic cells detected by FACS (Figure 4A) by showing cleaved caspase 3 induction in total protein extracts from control cells exposed to UV (no DOX). The absence of cleaved caspase 3 signal from ΔDboxGeminin-expressing cells confirmed that the discrete sub-G1 cell population observed upon ΔDboxGeminin induction (Figure 4A, lower left panel) does not represent apoptotic cells. As control, a short incubation with hydroxyurea (HU), to block DNA replication, did not induce any detectable apoptosis in cells synchronized for 20 hours. **C.** ΔDboxGeminin expression induction does not induce p53 activation (phosphorylation at serine 15), but does not inhibit its activation after UV-induced DNA damage, showing that ΔDboxGeminin does not prevent p53 induction. **D.** After ΔDboxGeminin expression induction (lower panels, +DOX) or not (upper panels, CTRL), mouse embryonic stem (ES) cells were fixed, incubated with an antibody against kinetochores (CREST) and then stained with DAPI (DNA). The number of CREST-positive dots per nucleus (indicated in the images) was determined with the IMARIS software after 3D image reconstruction. **E.** Number of CREST dots per nucleus in mouse ES cells in which ΔDboxGeminin expression was induced (red bars) or not (blue bars; control). Black dashed lines highlight the theoretical number of kinetochores per nucleus in 2C and 1C cells. The blue dashed line represents the average number of CREST dots observed in control cells to visualize the approach bias. The red dashed line indicates the average number of CREST dots observed in cells with fewer than 20 dots. With this approach, 20% of kinetochores were not detected in control cells. In ΔDboxGeminin-expressing cells with fewer than 20 dots, the average number of CREST dots was 15.7. This value would correspond to 20 kinetochores when the bias observed in control cells is taken into account. **F.** *The 1C cell population can re-enter the cell cycle after CDT1 overexpression*. At T0, ΔDboxGeminin expression was induced (+DOX) in asynchronous 3T3 cells for 24h and then DOX was washed off and cells were transfected with a CDT1-encoding plasmid or empty vector (T1). Cells were incubated with BrdU for 15 min just before fixation at T2 (left panels), or for 24 hours before analysis at T2 (right panels). During the last 24 hours, 15.3% of 1C cells transfected with CDT1 incorporated BrdU compared with 4.9% of control cells (empty vector). Upon CDT1 overexpression, the genome was actively replicated in 8.3% of 1C cells (BrdU^+^) during the 15 minutes prior to fixation, compared with 2.7% of control cells (empty vector). For each condition, the percentage of BrdU^+^ (red) and BrdU^−^ (blue) cells in the sub-G1 population is shown. The same experiment with non-induced (-DOX) cells is shown in Supplementary Figure 9.

To determine the number of chromosomes in cells arising from cell division of unlicensed cells, we induced ΔDboxGeminin expression and counted the number of kinetochores stained by the CREST polyclonal antibody in mouse embryonic stem cells that exhibit a stable diploid karyotype. We found an average of 31.5 dots per nucleus in control diploid cells (40 dots, one per chromosome, are theoretically expected) (see Figure 4D and 4E). This discrepancy could be explained by the close proximity of some kinetochores in interphase. In ΔDboxGeminin-expressing cells (but never in control cells), 7% of nuclei had up to 20 dots (mean: 15.7 dots), a proportion similar to the fraction of 1C cells detected by flow cytometry (see Figure 4A and 4E). The mean number of 15.7 dots per nucleus corresponds, when considering a similar bias of detection as in control cells, to 19.6 kinetochores, a number close to the theoretical 20 dots per nucleus in 1C cells.

### The 1C cell population can re-enter S phase

Finally, we investigated whether 1C cells could re-enter the cell cycle after stopping ΔDboxGeminin expression. We removed DOX 24 hours after induction to stop ΔDboxGeminin induction (T1, Figure 4F; control cells shown in Supplementary Figure 9) and transfected these cells with a CDT1-encoding plasmid (or empty vector; control) to counteract the effect of the remaining ΔDboxGeminin. 24h after CDT1 transfection (T2), 15.3% of 1C cells were BrdU-positive (Figure 4F, BrdU 24h, right panels) compared with 4.9% of 1C cells transfected with empty vector.

To rule out the possibility that these BrdU-positive 1C cells corresponded to diploid cells that replicated and died during the 24 hours preceding cell harvesting, we performed a short BrdU pulse (Figure 4F, BrdU pulse, T2+CDT1 and T2+empty vector) just before cell collection at T2 and confirmed that 1C cells can re-enter the S phase after re-establishment of origin licensing by CDT1 expression.

The generation of gametes by meiosis is the natural example of haploid cell formation characterized by the lack of DNA replication between meiosis I and meiosis II ^32^. Interestingly, we observed that geminin expression was strongly upregulated during testis gametogenesis, particularly when meiosis is induced (Supplementary Figure 10). Therefore, the stable geminin-induced “reductive somatic mitosis” reported here could mimic some events occurring during the formation of haploid cells during development.

## DISCUSSION

Here, we report that somatic mammalian cells can enter mitosis without DNA replication. We found that in normal adult mouse somatic cells, the different checkpoint response pathways are operational, but they are not induced when DNA replication initiation is fully prevented by complete licensing inhibition. We cannot exclude the existence of a still unknown checkpoint that might detect a defect at an earlier step of pre-RC formation, at the level of ORC or CDC6 for example. However, our data reveals that the absence of replication licensing (i.e. defined as the loading of the MCM helicase onto DNA replication origin) is not sensed by the cell cycle machinery. Importantly, the mitotic entry of cells with an unreplicated genome triggered by licensing inhibition does not rely on cell synchronization^25^(Supplementary Figure 6).

The majority of non-replicated cells undergo cell cycle arrest and senescence due to mitotic failure (Supplementary Figure 11). This is reminiscent of the senescence observed in normal cells after interference with mitotic progression^29,33,34,35^. Mitotic delay or failure can induce γH2AX and the DNA damage response in normal diploid cells that are undergoing tetraploidization or mitotic perturbations^36,28^. Here, we showed that mitotic entry precedes p53/p21 activation in unlicensed cells. Therefore, DNA damage (and the subsequent cell cycle block in G1) appears to be the consequence of mitotic failure of unlicensed cells rather than of the absence of licensing or DNA replication. This suggests that mitotic failure induces senescence regardless of any replicative process and explains why cancer cells, which are more susceptible to cell death upon mitotic perturbations than normal cells, massively die upon licensing inhibition^36,37,30^.

A discrete part of unlicensed cells can establish a mitotic spindle and divide despite the absence of DNA replication. During these “reductive somatic mitoses” induced by ΔDboxGeminin, the absence of replicated sister chromatids should prevent the spindle assembly checkpoint fulfilment and, therefore, anaphase onset. However, it has been reported that the spindle assembly checkpoint can be satisfied after prolonged mitosis ^38,39,40^. Similarly, we observed an increase of mitosis duration in unlicensed cells. Therefore, a discrete population of unlicensed somatic cells can ultimately complete mitosis, giving rise to two daughter cells with 1C DNA content that are viable and show a relative homogenous DNA content, with part of them containing half genome (20 chromosomes). These results suggest the possibility to develop new experimental approaches to generate haploid cells from a wide range of somatic mammalian cells.

## Supporting information

related to Figure 3C minus DOX

related to Figure 3C plus DOX

related to figure 3B_plus DOX

## FIGURE AND SUPPLEMENTARY FIGURES LEGENDS

Each figure and Supplementary Figure legend is presented on the same page as its figure.

## SUPPLEMENTARY MOVIE LEGENDS

**Sup movie 1. Time lapse related to Fig 3B**. 3T3 cells expressing H2B-GFP were induced for ΔDboxGeminin. Images were acquired every 15 min. H2B-GFP is shown in green and overlaid with Bright field images.

**Sup movie 2. Time lapse related to Fig 3C** showing non-induced 3T3 cells expressing H2B-GFP and Tag-RFP-a-tubulin. Images were acquired every 20 min. H2B-GFP is shown in green, Tag-RFP-a-tubulin in Red and overlaid with Bright field images. 4 frames per second are displayed.

**Sup movie 3. Time lapse related to Fig 3C**. 3T3 cells expressing H2B-GFP and Tag-RFP-a-tubulin were induced for ΔDboxGeminin. Images were acquired every 20 min. H2B-GFP is shown in green, Tag-RFP-a-tubulin in Red. 1 frame per second are displayed.

## MATERIAL and METHODS

### Plasmids and constructs

Deletion of the destruction box (Dbox) and addition of N-terminal Flag-HA epitopes were performed by PCR. ΔDboxGeminin was finally cloned in the pTRIPZ vector using the *Age*1-*Mlu*1 cloning sites (Supplementary Information 1). The plasmid encoding the shRNA against CDT1 was from Sigma-Aldrich (TRCN0000174484). The CDT1-pCDNA3 plasmid is described in Coulombe *et al.*^42^.

### Preparation of viral particles and infection

HIV-derived vectors pseudo-typed with the VSV-G envelope protein were produced by transient co-transfection of the Gag-Pol packaging construct psPAX2, the envelop plasmid pMD.2G and the self-inactivating (SIN) HIV-1–derived vector coding for the ΔDboxGeminin or shRNA of interest. 36 hours after transfection, particles were harvested and filtered before infection of different cell lines.

### Cells and media

Mouse embryonic fibroblasts (MEFs) were derived as previously described ^43^. Briefly, 13.5-day-post-coitum wild type C57BL6 mouse embryos were dissected and gonads, internal organs and head were removed before MEF isolation. Tissues were then dissociated with 0.05% trypsin/EDTA (Gibco, Invitrogen) at 37°C for 15 minutes to isolate single cells. To obtain tail tip fibroblasts (TTFs), tail tips from adult mice were sliced into small pieces, trypsinised and plated to derive fibroblast cultures. All fibroblasts were expanded for three passages in Dulbecco’s Modified Eagle Medium (DMEM) (Invitrogen) with 10% foetal bovine serum (FBS) (catalogue no. S1810; Biowest) before being used for experiments.

NIH 3T3, TTF, MEFs and T98G cells were grown in high-glucose DMEM (Invitrogen) supplemented with 10% FBS, 2 mM L-glutamine (Invitrogen) and 1 mM sodium pyruvate (Sigma). HCT116 p53^+/+^ cells were grown in McCoy’s 5a medium modified with 10% FBS. The mouse ES cell line CGR8 was from C. Crozet (Institute of Human Genetics, Montpellier, France) and was cultured as described in Ganier *et al.* ^43^. Specifically, ES cells were grown on 0.1% gelatine without feeders in ES cell medium [Glasgow minimum essential medium (Invitrogen) with 10% FBS, 0.1 mM β-mercaptoethanol, 1mM sodium pyruvate, 1% nonessential amino acids (Gibco), 2 mM L-glutamine and 1,000 U/mL leukaemia inhibitory factor (LIF) (ES-GRO)] at 37 °C in 5% CO_2_.

To obtain stable cell lines that express inducible ΔDboxGeminin, exponentially growing cells were infected with HIV-derived viral particles that were previously filtered and concentrated by ultracentrifugation. 48 hours after infection, cells were selected with 2 μg.mL^−1^ puromycin. For ES cells, clonal selection was performed to ensure homogeneous ΔDboxGeminin expression upon doxycycline (DOX) induction.

Primary TTFs and MEFs were infected at sub-confluence with viral particles containing pTRIPZ-ΔDboxGeminin and processed immediately for experiments.

Expression of ΔDboxGeminin was induced by adding 1-2 μg.mL^−1^ DOX, (Clontech) in the medium.

### Immunofluorescence

Cells were plated on coverslips, washed twice in PBS and then fixed in 3% paraformaldehyde (PFA) at room temperature (RT) for 10 min, washed with PBS, and permeabilized with PBS/0.2% Triton X-100 for 5min except for CREST (abcam, ab227535) and α-tubulin (Sigma, DM1A T6199) staining for which cells were fixed in cold methanol for 5 min. Then, cells were washed three times in PBS/2% BSA for 10min, incubated with primary antibodies diluted in blocking buffer (PBS/0.5% Tween/2% BSA) at 4°C overnight, except for the CREST antibody (diluted in PBS/0.5% Tween/5% non-fat milk and added at RT for 1 hour). Primary antibodies used are anti-^ser10^phosphoH3 (Ozyme, 9701S), anti-^ser139^γH2AX (Millipore, 05-636) and anti-nucleolin (Novus, NB100-2239). After, three washes in PBS/0.5% Tween, cells were incubated with secondary antibodies diluted in blocking buffer for 1 hr and DNA stained with Hoechst. Cells were mounted on glass slides with Prolong (Sigma-Aldrich). For co-staining with EdU, EdU was detected prior to immune-detection according to the manufacturer’s instructions.

Cells were observed using wide-field fluorescence microscopes (Leica, Germany and Carl Zeiss, Germany) and 63x 1.4NA PL APO lenses. Filter sets allowing the observation of quadruple staining were used (Carl Zeiss FS49, FS38HE, FS43 and FS50). Images were acquired using a Hamamatsu ORCA Flash4 sCMOS camera. For chromosomes counting using CREST staining, ES cells were fixed and stained as described above for EdU and CREST staining 24 hrs after ΔDboxGeminin induction. Images were acquired with a Zeiss Axioimager Z3 Apotome and stacks were acquired according to the Nyquist criterion. For better contrast, a Grid Projection Illumination Microscopy (aka Apotome) was used. 3D reconstruction was performed using the Imaris software (Bitplane, Oxford Instruments). Briefly the nuclear envelope was modelled using isosurface detection, and CREST foci were detected using spot detection. The number of spots/envelope was automatically derived using the split into surface object Xtension.

3D-SIM imaging was performed using an OMX-V3 microscope (General Electrics) equipped with 405 nm, 488 nm and 561 nm lasers and the corresponding dichroic and filter sets. The Far Red channel, used to detect EdU incorporation, was acquired in wide-field mode only. Reconstruction and alignment of the 3D-SIM images was carried out with softWoRx v 5.0 (General Electrics). 100nm green fluorescent beads (Life Technologies) were used to measure the optical transfer function (otf) for the 405 and 488 channels, and 100 nm red fluorescent beads (Life Technologies) were used to measure the otf for the 561 channel. 170 nm TetraSpeck beads (Life Technologies) were employed to measure the offsets and rotation parameters used in the image registration. Reconstructed 3D-SIM images were analysed with the Image J 1.4.7v software.

### Metaphase spreads

pTRIPZ-ΔDboxGeminin-infected MEFs were synchronized in G0 by confluence and then released by splitting them in fresh medium containing 15% FBS + 10 μM EdU with or without DOX. 15 hours later, 100 nM Karyomax was added in the medium for 7 hours. Then, metaphase spreads were prepared as described in Eot-Houllier *et al.* 2008 ^44^, fixed and processed for EdU detection and DNA staining.

### Video-microscopy

For video microscopy experiments, ΔDboxGeminin-expressing 3T3 cells were infected with viruses encoding H2B-GFP and processed for time lapse experiments without selection. 3T3 cells were seeded on μ-Dish 35 mm, high Grid-50 Glass Bottom gridded coverslips and were synchronized by serum starvation, as described above, and released in the presence of 10 μM EdU and DOX or not. Tilescan Timelapse images were acquired every 5 minutes for 20 hrs. To ensure a large sample size, the four whole quadrants of the Ibidi dish were recorded, then fixed and processed for EdU detection, as described above.

### Flow cytometry

For propidium iodide (PI), BrdU/PI and ^ser10^PH3/PI staining, cells were fixed in cold 70% ethanol/PBS and then processed for indirect immunofluorescence, as described above. To detect cyclin B1/PI, MPM2/PI and FLAG/PI, cells were fixed in 3% PFA in PBS at RT for 15 min, permeabilized in 0.5% Triton X-100 for 5 min and processed for indirect immunofluorescence: anti-cyclin B1 (Santa Cruz, sc-245), anti MPM2 (Millipore, 05-368), anti-FLAG (Sigma M2, F1804), anti-BrdU (BD Biosciences, B44). For DNA staining, cells were treated with 50 μg.ml^−1^ RNase A (Sigma, R6513) and stained with 25 μg.ml^−1^ PI (Sigma, P4864).

For BrdU detection, cells were incubated for the indicated times before harvesting and fixation. Then, cells were incubated with 2 N HCl for 30 min and washed in PBS 0.2% Tween (PBS-T) with 5% BSA before BrdU detection.

For EdU/^ser10^PH3/DRAQ5 staining, fixed cells were first processed for EdU detection following the manufacturer’s instructions, and then incubated with the anti-^ser10^PH3 antibody in PBS-T/5% BSA at 4°C overnight. Cells were then washed twice in PBS-T and incubated at RT with the anti-rabbit PE antibody (Thermofisher) (1/200 in PBS-T/5% BSA) and after two washes in PBS-T, stained with 2.5 μm DRAQ5 (DNA) and 50 μg.ml^−1^ RNase A at RT for 1hr.

Cells were analysed with a FACSCalibur flow cytometer using the CellQuestPro software and a Miltenyi MACS quant cytometer.

### Cell synchronization and treatments

To synchronize 3T3 cells in G0/G1, cells were incubated in serum-depleted medium (0.5% FBS) for 36 hrs. Cells were then released in medium supplemented with 15% FBS +/− DOX and +/− 10 μM EdU. Primary TTFs and MEFs were synchronized by confluence 16 hrs after infection and released by splitting (1:4 ratio).

To express CDT1, 3T3 cells were transfected with 5 μg of the pCDNA3-CDT1 plasmid or empty vector using Nucleofector™ II, Amaxa Biosystem, AAD-1001N. To induce DNA damage, synchronized cells were exposed to UV (0.002 joules) using the Bioblock Scientific BLX-254 system upon release, or incubated with 2mM hydroxyurea (Sigma, H8627) for 20 hrs. To block mitotic progression, 9μM RO-3306 (Sigma, SML0569) was added in the medium upon G0/G1 release.

### Cell proliferation curves

At day 0, 10^3^ cells (3T3, T98G, HCT116) were seeded at sub-confluence and then harvested every day for cell counting. Values represent the mean ± SD of three experiments.

### Senescence-associated β-galactosidase staining

Sub-confluent asynchronous mouse TTFs were infected with ΔDboxGeminin-encoding viral particles and cultured in the presence, or not of DOX for 14 days. Senescence-associated β-galactosidase (SA-β-gal) activity was assessed as described ^45^. Briefly, cells were washed twice in PBS, fixed in 2% formaldehyde/0.2% glutaraldehyde at RT for 5min, washed twice with PBS and incubated at 37°C in freshly prepared SA-β-gal solution (1 mg.ml^−1^ of X-gal; 40 mM citric acid; sodium phosphate, pH 6.0; 5 mM K_4_[Fe(CN)_6_] 3H_2_O, 5 mM K_3_[Fe(CN)_6_]; 150 mM NaCl_2_ and 2 mM MgCl_2_) overnight. Cells were washed with PBS, fixed with methanol and air-dried before analysis.

### Western blotting

To prepare total cell extracts, cells were harvested, washed twice with PBS, lysed in 2x Laemmli buffer, sonicated (3×30 seconds) and boiled for 5min. To obtain chromatin-enriched fractions, a protocol adapted from Lutzmann et *al.* ^46^ was used. Briefly, cells were harvested, washed with PBS and lysed on ice in CSK buffer [150 mM NaCl, 10 mM HEPES pH 7.5, 300 mM Sucrose, 1 mM MgCl_2_, 1 mM EDTA, 1 mM ATP⋅MgCl_2_, 1 mM DTT, 0.5% Triton X-100, phosphatase-inhibitors (Calbiochem), the protease inhibitors leupeptin, aprotinin, and pepstatin at a final concentration of 10 μg/ml] for 30min. Lysed cells were then centrifuged at 3800g, 4°C, for 5 min to obtain the soluble fraction. Pellets were washed twice with CSK buffer on ice for 5 min, centrifuged at 3800g, 4°C for 5min, solubilized in 2× Laemmli Sample Buffer, then sonicated (3×30s) and boiled for 5min. Extracts were then analysed by SDS/PAGE and western blots were performed using the following antibodies: anti-^ser10^phospho-H3 (Ozyme; 9701), , anti-histone H3 (ab1791; Abcam), anti-p53 (Cell Signalling, 2524), anti-^ser15^-phospho-p53 (Ozyme, 12571), anti-GAPDH (abcam, ab9484), anti-MCM2 (Santa Cruz, sc9839), anti-MCM4 (ref), anti-cyclin A2 (Santa Cruz, sc-533229), anti-CDC6 (Santa Cruz, sc-6316), anti-Geminin (Santa Cruz, sc8448 and sc-74496), anti-p21 (Santa Cruz, sc-397), anti-phospho-RB (Cell signalling, 9307), anti-RPA32 (abcam, ab2175), anti-phosphoRPA32 (bethyl laboratories, A300-245A), anti-Plk1 (abcam, ab17056), anti-CDT1 (gift from M. Fujita), anti-histone H4 (Millipore, 07108), anti-cleaved Caspase 3 (Cell Signalling, 9661), anti-PCNA (Sigma, P8825), anti-p27 (Santa Cruz, sc-528), anti-53BP1 (Millipore, MAB-3804), anti-CHK2 (Millipore, 05-649), anti-phospho-CHK1 (Cell Signalling, 2341), anti-^Thr2609-^phospho-DNA-PK (Thermo Fisher), anti-Actin (Sigma, A4700), anti-SYCP3 (gift from P. de Boer).

### Testis extracts

Sv129/B6 mice were euthanized at the indicated ages and testis protein extracts were prepared in RIPA buffer (25 mM Tris, pH 7–8; 150 mM NaCl; 0.1% SDS; 0.5% sodium deoxycholate; 1% Triton X-100; the protease inhibitors leupeptin, aprotinin and pepstatin at a final concentration of 10 μg/ml).

## Author Contributions

O.G. contributed to research design, performed and analysed the experiments, and co-wrote the manuscript. M.L. contributed to the analyses of the experiments and performed the experiments on mouse testes extracts. J.C. contributed to video-microscopy experiments and to the determination of chromosomes number in interphase ES cells. C.L. produced viral particles encoding ΔDboxGeminin and shRNA against CDT1. I.P. and S.Z. contributed to experiments on ES cells. C.T. contributed to the analysis of experiments. M.M. supervised the project, contributed to research design and to the analyses of the experimental results, and co-wrote the manuscript.

---

**Supplementary Figure 1:**
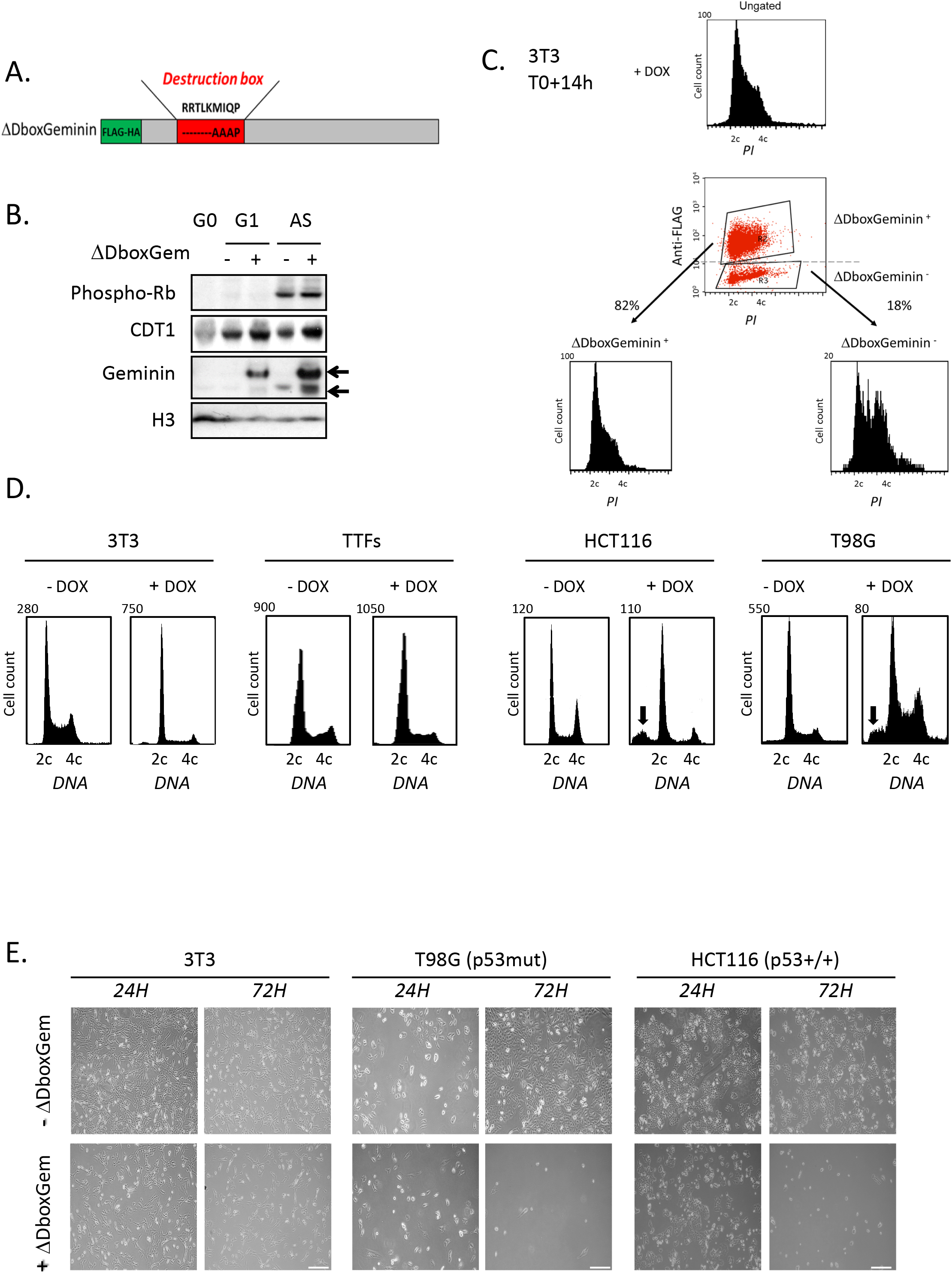
Expression of ΔDboxGeminin blocks DNA replication licensing and inhibits cell proliferation in untransformed somatic cells but triggers cell death in cancer cells. **A.** Schematic representation of the ΔDboxGeminin construct showing the amino acid sequence of the destruction box (Dbox) and the three amino acids replaced by alanine residues. **B.** *Deletion of the destruction box in geminin stabilizes its expression in G1*. Analysis by western blotting of total extracts from asynchronous (AS) 3T3 cells or synchronized in G0 or in G1 after induction or not (+ or −) of ΔDboxGeminin expression by incubation with doxycycline (DOX). The G1 phase status of cells was confirmed by CDT1 expression and the absence of phosphorylated pRB. In G1, endogenous geminin (lower band) is degraded, but not FLAG-HA-tagged ΔDboxGeminin (upper band). H3 was used as loading control. **C.** Heterogeneous expression of ΔDboxGeminin upon DOX induction. Cell cycle profile of 3T3 cells synchronized in G0 and then released in the presence of doxycycline (DOX) to induce ΔDboxGeminin expression for 14 hours. Cells were fixed, incubated with an anti-FLAG antibody (to detect ΔDboxGeminin-positive cells) and stained with propidium iodide (PI; DNA content) and analysed by flow cytometry (upper panel). Bi-parametric representation of DNA content (PI) and FLAG signal (middle panel). Different cell cycle profiles of ΔDboxGeminin-positive (left) and -negative (right) 3T3 cells (lower panels). **D.** Cell cycle profiles of the indicated cell lines 48 hours after or not induction of ΔDboxGeminin expression (+/− DOX) showing the spread profile of the sub-G1 population (black arrows) in the cancer cell lines (HCT116 and T98G), but not in non-transformed cells (adult Tail Tip Fibroblasts, TTFs, and mouse 3T3 fibroblasts). **E.** Representative phase-contrast images of 3T3, HCT116 and T98G cells 24 hours and 72 hours after induction (lower panels) or not (upper panels) of ΔDboxGeminin (+ or − DOX) showing cell death in cancer cells, but not in 3T3 fibroblasts. Scale bars = 200 μm.

**Supplementary Figure 2:**
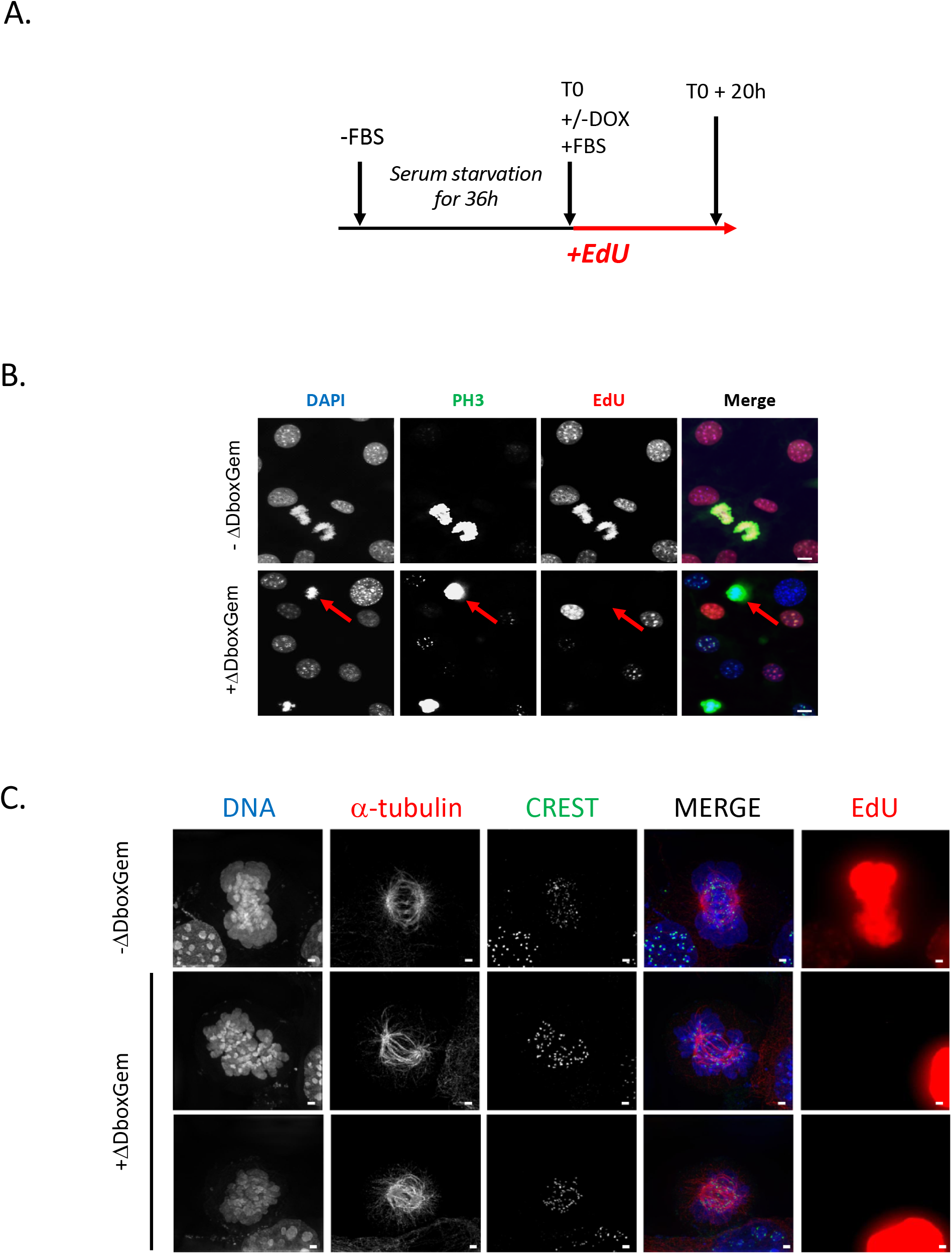
Expression of ΔDboxGeminin induces mitotic entry of unreplicated genome. **A.** *Expression of ΔDboxGeminin induces mitotic entry of EdU-negative cells*. 3T3 cells were synchronized in G0 and then released in the presence of EdU (+/− DOX to induce ΔDboxGeminin expression). At T0+20h, cells were fixed to assess EdU incorporation and stained with different antibodies. **B.** In control cultures (- DOX; no ΔDboxGeminin expression) all mitotic cells were EdU-positive (i.e. post-replicative cells). Conversely, in cells that express ΔDboxGeminin (+DOX), some unlicensed mitotic cells without detectable EdU staining were observed (red arrow). PH3, histone 3 phosphorylated on serine 10 (a mitosis marker). Scale bars = 5 μm. **C.** Super-resolution microscopy analysis of fixed 3T3 cells incubated with antibodies against α-tubulin and kinetochores (CREST). After release from G0, cells were labelled with EdU for 20 hours, in the presence or not of DOX to induce (+) or not (−) ΔDboxGeminin expression. Representative images of one control mitotic cell (upper panels) and two unlicensed mitotic cells (middle and lower panels) with bipolar spindles despite the absence of DNA replication. Scale bars = 1 μm.

**Supplementary Figure 3:**
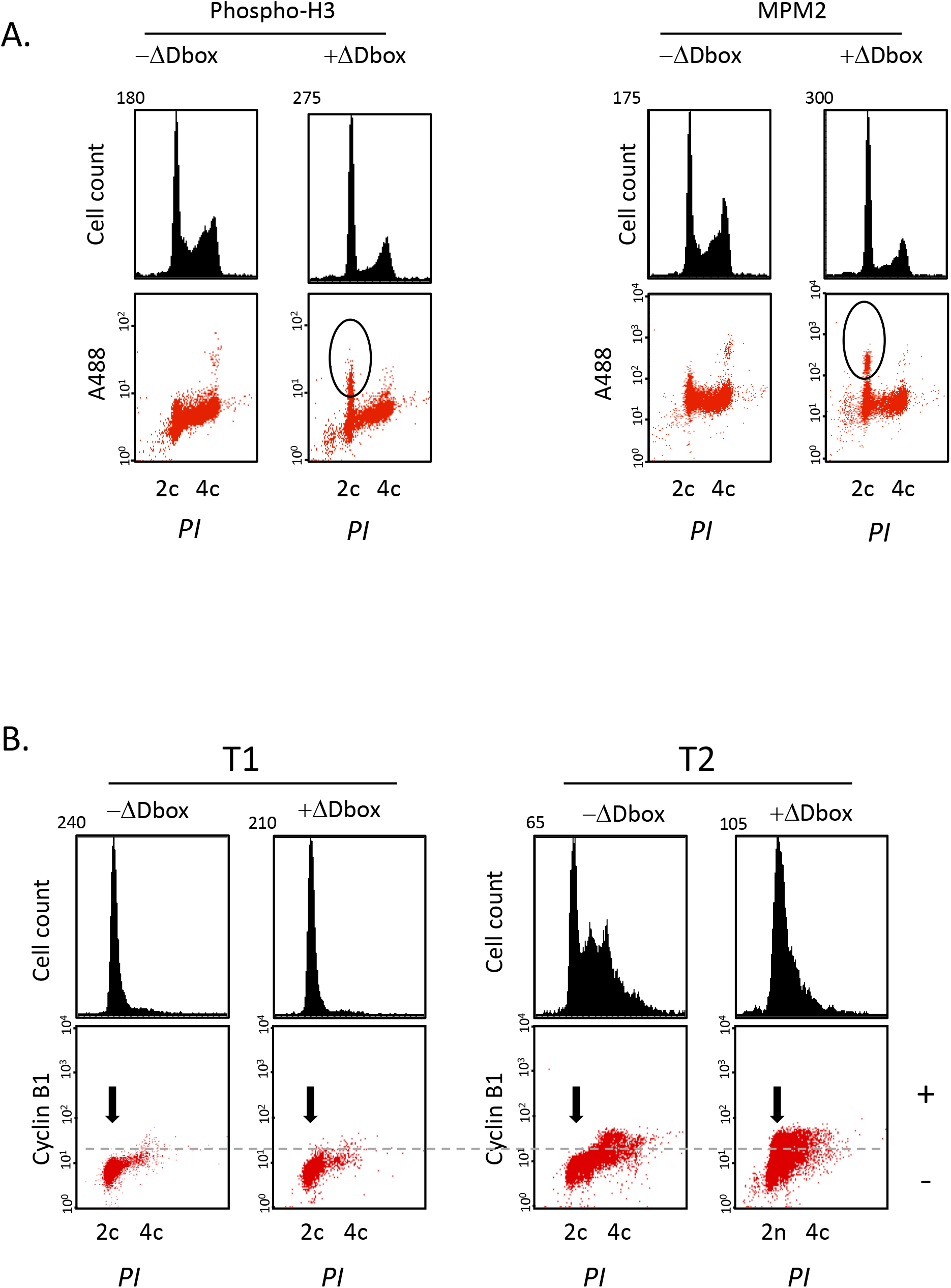
Expression of ΔDboxGeminin does not prevent the appearance of mitotic markers despite the absence of DNA replication. **A.** 3T3 cells were synchronized in G0 and released with or without (+ and – DOX) induction of ΔDboxGeminin expression for 20 hours. Cells were then fixed and the expression of ^ser10^PH3 and MPM2 (a mitosis-specific phosphorylated epitope) was analysed by flow cytometry to confirm the presence of mitotic cells with 2C DNA content (propidium iodide; PI) after induction of ΔDboxGeminin expression (highlighted by black circles). **B.** Cyclin B1 and DNA content were analysed by flow cytometry in synchronized 3T3 cells at T1 and T2 (7 hours and 17 hours post-release and ΔDboxGeminin induction, respectively). Similarly to ^ser10^PH3 and MPM2, cyclin B1 accumulated in 2C cells (black arrow) at T2 after ΔDboxGeminin induction.

**Supplementary Figure 4:**
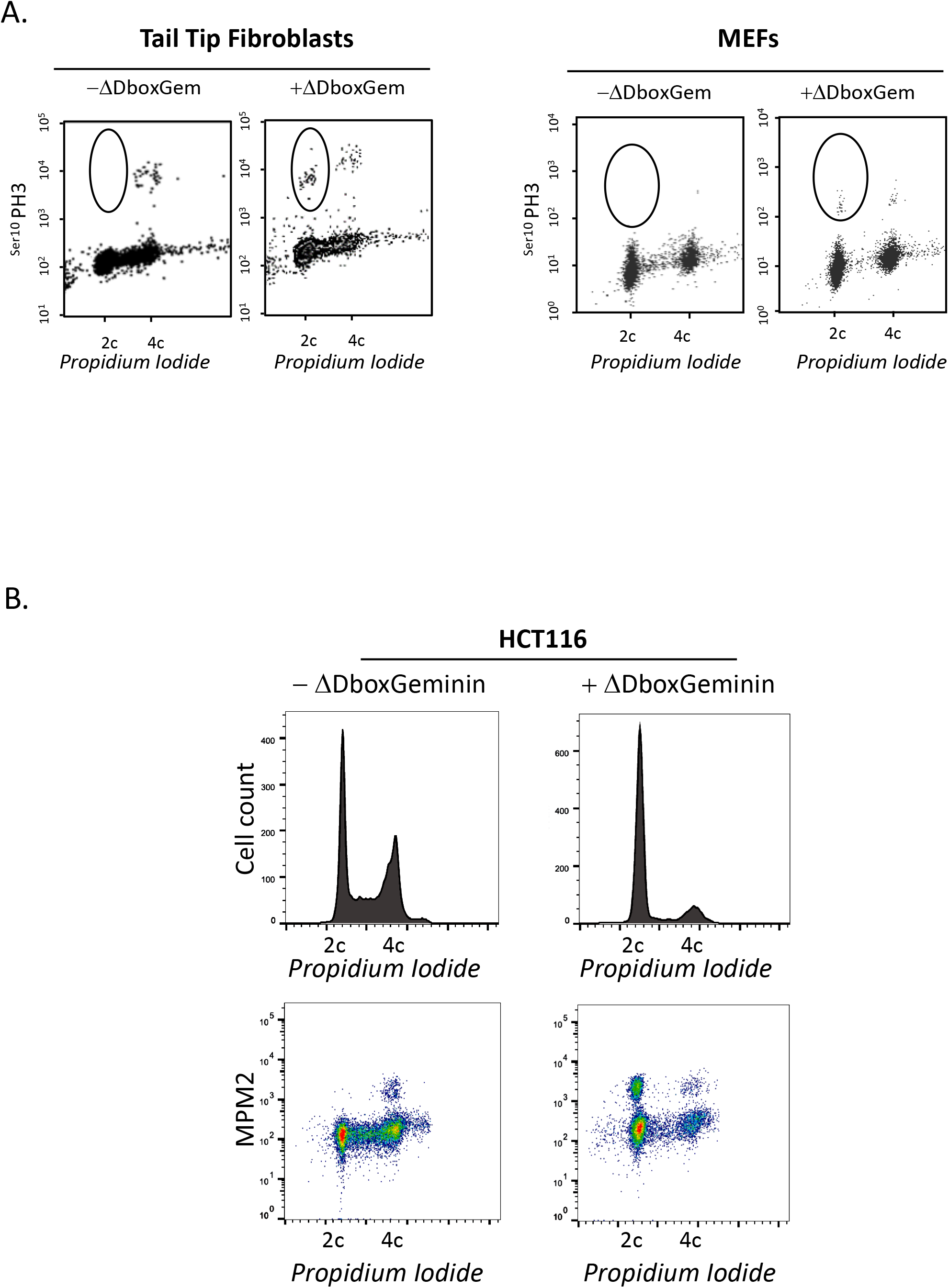
Mitotic entry of unlicensed cells occurs in both normal and cancer cells. **A.** Mouse adult tail-tip fibroblasts (left panels) and embryonic fibroblasts (MEFs) (right panels) were infected with ΔDboxGeminin-encoding viruses, synchronized in G0 and released in the cell cycle. The DNA content and ^ser10^PH3 level were measured by flow cytometry 20 hours after release and induction or not of ΔDboxGeminin expression (+ and – ΔDboxGem). Black circles highlight the ^ser10^PH3-positive 2C cell populations. **B.** HCT116 cancer cells were incubated with 2 mM thymidine for 24 hours to synchronize them in G1/S. Cells were then released in the presence (+DOX) or absence (-DOX) of doxycycline for 24 hours to induce or not ΔDboxGeminin expression. Cells were then fixed and the presence of MPM2 (a mitosis-specific phosphorylated epitope) was analysed by flow cytometry to confirm the presence of mitotic cells with 2C DNA content (propidium iodide; PI). The upper panels show the cell cycle profiles (propidium iodide), while the lower panels show the MPM2 signal in cells with different DNA content (bi-parametric analyses).

**Supplementary Figure 5:**
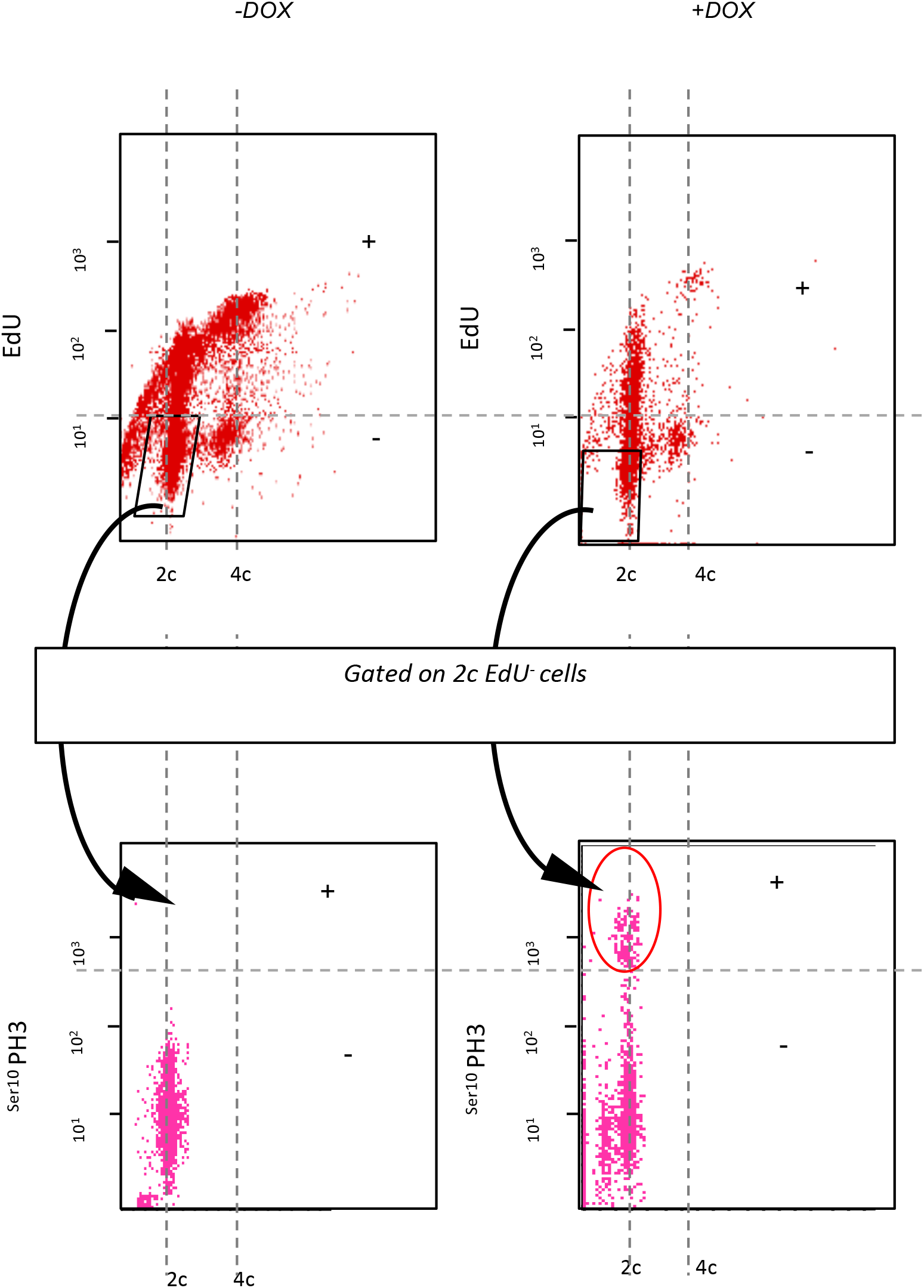
Simultaneous three-color FACS analyses of 3T3 cells in which ΔDboxGeminin expression was induced (right panels, +DOX) or not (left panels, -DOX). Bi-parametric representation of EdU detection in 2C and 4C cells. EdU-negative 2C cells were then gated to assess ^ser10^PH3 staining. The circle highlights the EdU-negative and ^ser10^PH3-positive 2C cell population observed only upon ΔDboxGeminin expression induction.

**Supplementary Figure 6:**
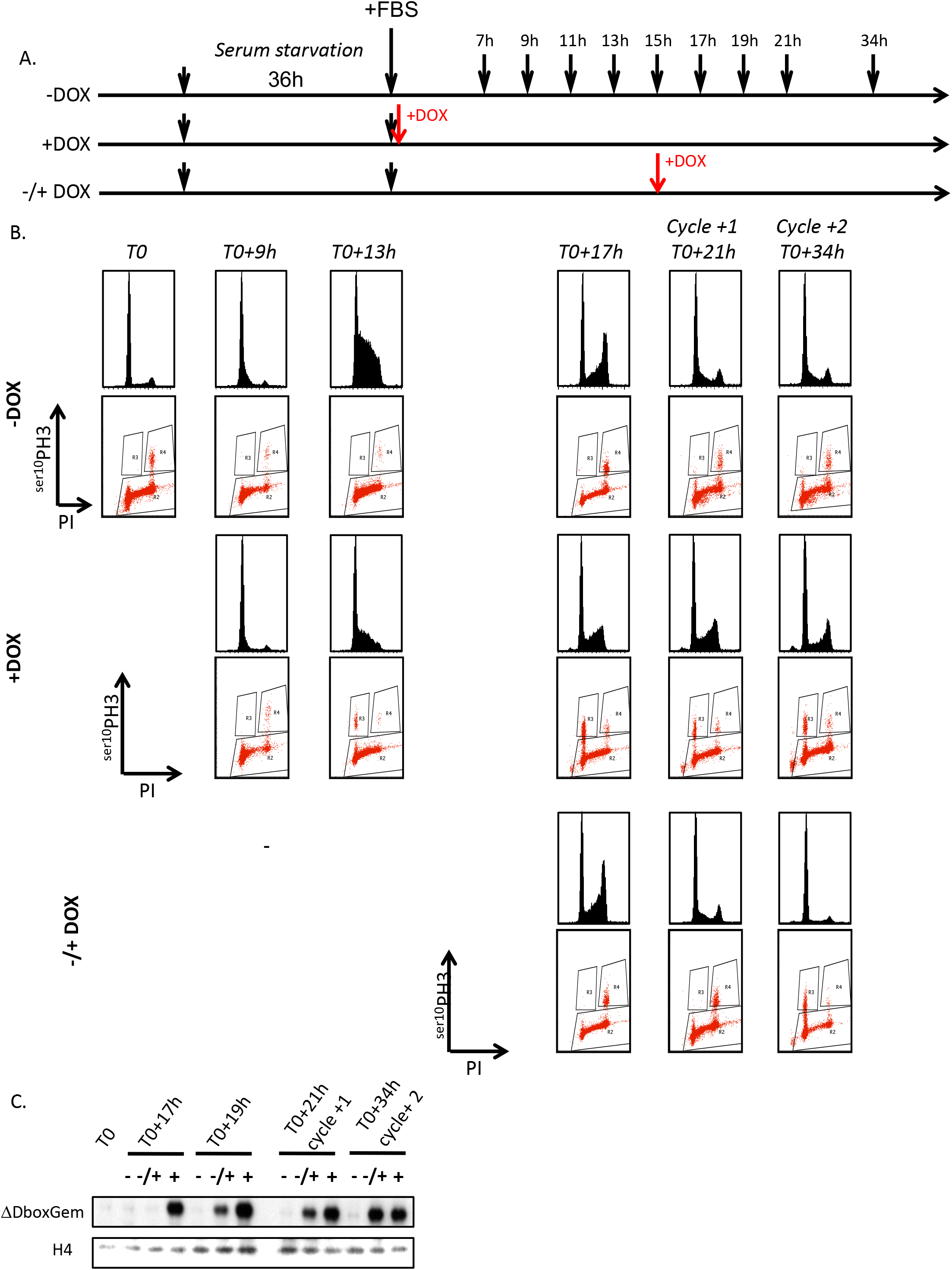
ΔDboxGeminin effect occurs in G1 and does not rely on cell cycle re-entry from quiescence. **A.** Scheme of the experiment in which ΔDboxGeminin expression was induced either upon release from quiescence (+DOX) or after licensing (−/+DOX), or never induced (- DOX). At different time points upon release from G0, cells were fixed and ^ser10^PH3 level relative to the DNA content (propidium iodide staining) was analysed by flow cytometry. FBS, foetal bovine serum. **B.** *Only ΔDboxGeminin expression induction in G1 leads to mitotic entry of cells with 2C DNA content*. For each condition (-DOX, +DOX and −/+ DOX), the upper panels show the cell cycle profiles, while the lower panels show the ^ser10^PH3 signal in cells with different DNA content (bi-parametric analyses). **C.** Western blots showing the expression of inducible ΔDboxGeminin in the indicated conditions. Non-degradable ΔDboxGeminin can be detected at T0 + 19h when the induction is performed at 17h post-release. In this context, the 2C population starts to appear when control cells enter the second round of mitosis.

**Supplementary Figure 7:**
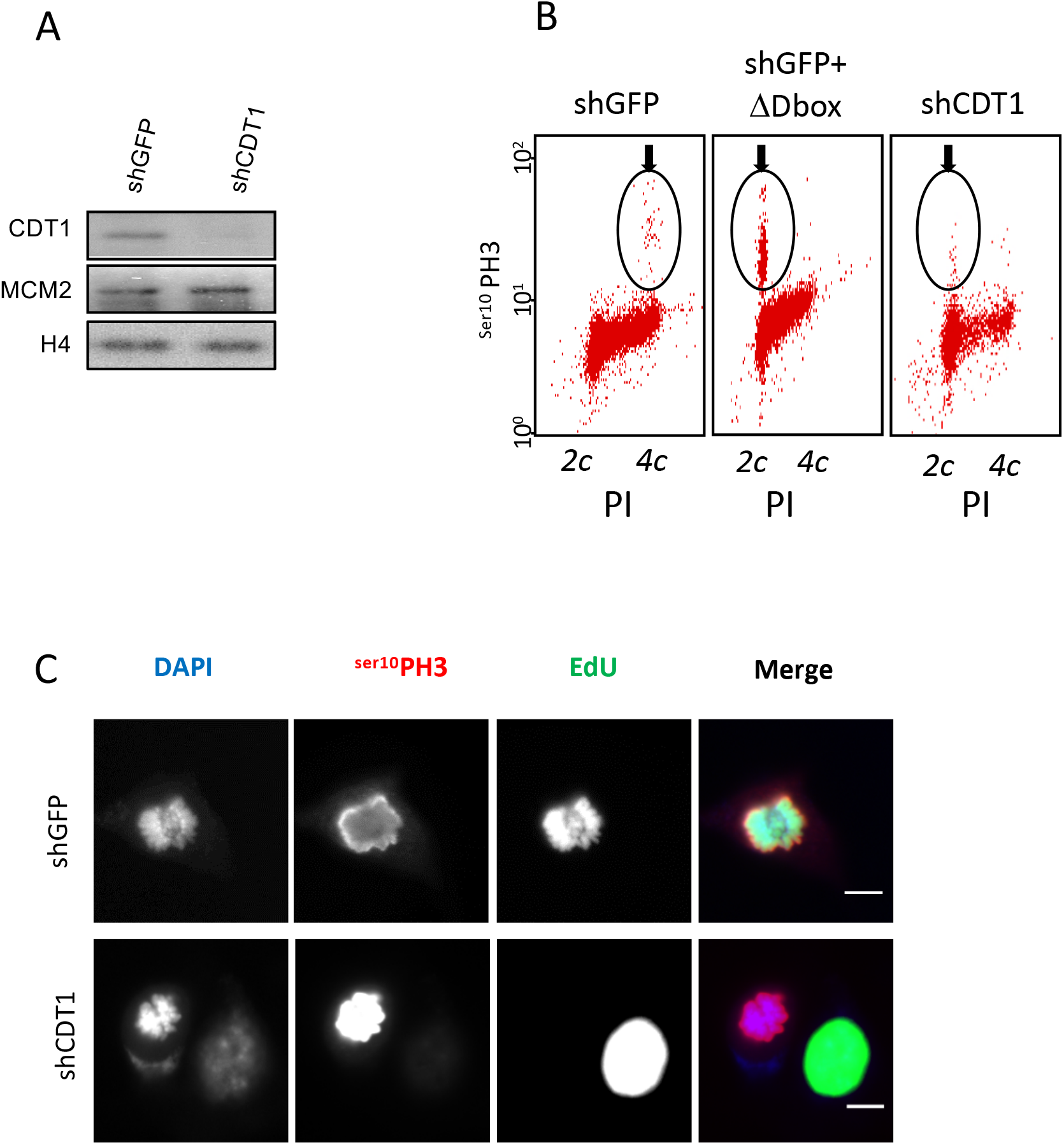
CDT1 downregulation mimics the effects of ΔDboxGeminin expression. **A.** 3T3 cells were infected with plasmids encoding shRNAs against GFP (control) or CDT1 (shCDT1), synchronized in G0, and then released in the cell cycle. Total protein extracts were analysed by western blotting to assess the efficiency of CDT1 silencing. **B.** At 20 hours post-release, cells were fixed and ^ser10^PH3 level and DNA content (propidium iodide, PI, staining) were analysed by flow cytometry. **C.** To confirm the mitotic status of non-replicated shCDT1-expressing 3T3 cells, cells were synchronized in G0, released in the presence of EdU for 20 hours, fixed and incubated with an anti-^ser10^PH3 antibody to detect by immunofluorescence the presence of EdU-negative cells in the G2/M phase (=unlicensed mitotic cells, arrows). Scale bars = 5 μm.

**Supplementary Figure 8.**
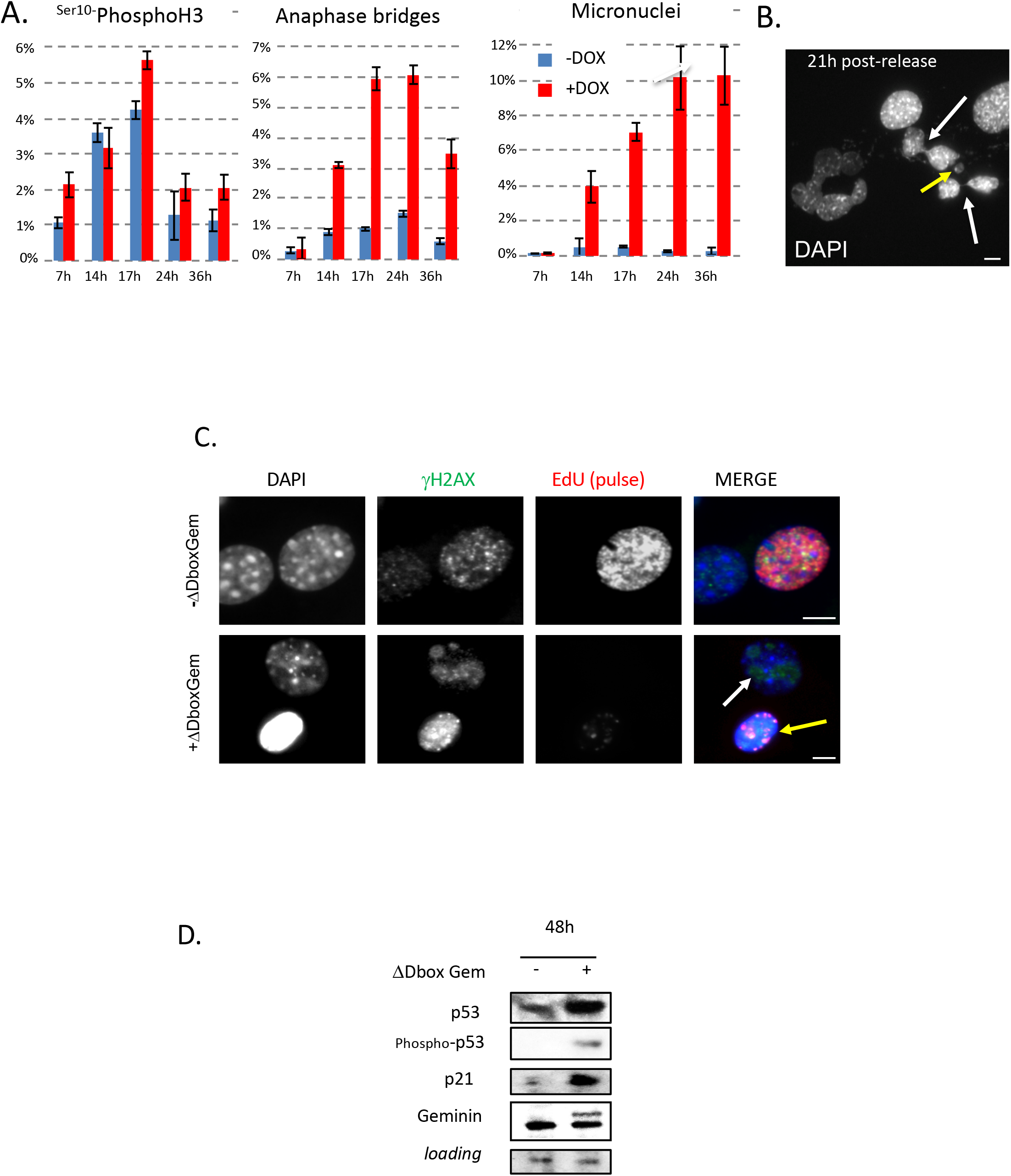
Mitotic failure of unlicensed cells induced anaphase bridges and nucleolar DNA damages. **A.** At different time points after release from G0, and incubation (+DOX, red bars) or not (-DOX, blue bars) with DOX to induce ΔDboxGeminin expression, 3T3 cells were fixed and stained with DAPI (to determine the proportion of anaphase bridges and micronuclei) and with an anti-^ser10^PH3 antibody (to determine the number of mitotic cells by immunofluorescence). Anaphase bridges and micronuclei started to appear after the first wave of mitoses (from 14h post-release), suggesting that they might result from mitotic failure. Data are the mean ± SEM of nine fields of view from three experiments). **B.** Expression of ΔDboxGeminin in 3T3 cells induces anaphase bridges (white arrows) and micronuclei (yellow arrow) after the first mitotic wave (DAPI staining at 21h after release from G0). Scale bars = 5 μm. **C.** 3T3 cells were synchronized in G0 and then incubated with or without doxycycline (+/−ΔDboxGeminin) for 24 hours. Then, cells were incubated with EdU for 15 min, fixed and stained for γH2AX, EdU and DAPI. In some ΔDboxGeminin-expressing cells, γH2AX staining is observed, mainly restricted to DAPI-negative nuclear structures, suggesting the nucleolar localization of damaged DNA. None of these cells could replicate, as indicated by the absence of EdU incorporation. The nucleolar γH2AX signal was weaker and differently distributed in ΔDboxGeminin-expressing cells (white arrow) compared with replicating control cells or cells that do not efficiently express ΔDboxGeminin upon induction (yellow arrow) and therefore show canonical DNA damage induced by basal replicative stress (EdU-positive). Scale bars = 5 μm. **D.** 48 hours after release from G0 and incubation with (+) or without (−) DOX, total protein extracts by western blotting. Upon ΔDboxGeminin expression (+DOX), p53 and p21 were induced, as previously described following the same timing protocol ^41^.

**Supplementary Figure 9:**
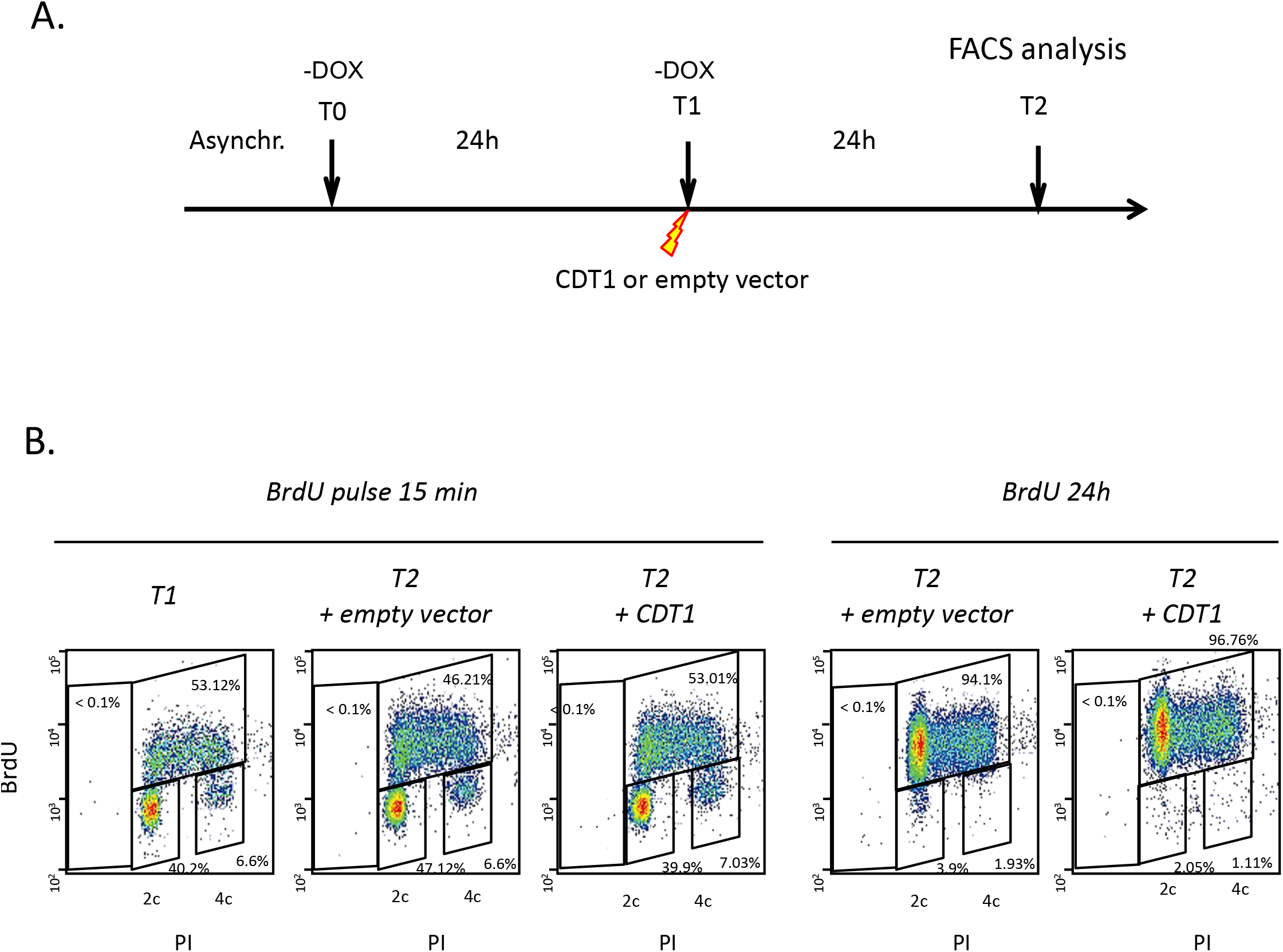
Cell cycle and BrdU incorporation in non-induced cells transfected with the CDT1-encoding plasmid or empty vector, related to Figure 4F. **A.** Scheme of the experiment related to Figure 4F for control cells in which ΔDboxGeminin expression was not induced (-DOX) at T0. **B.** Bi-parametric analysis of DNA content (propidium iodide; PI) and BrdU incorporation in non-induced 3T3 cells that were transfected with empty vector or the CDT1-encoding plasmid at T1 and incubated with BrdU for 15 minutes prior to harvesting at T2 (to visualize replicating cells, 24 hours after transfection) or for 24 hours (to quantify the proportion of cells that have replicated DNA during the last 24 hours). Cells were fixed and stained for DNA (propidium iodide, PI), and BrdU incorporation assessed. Results show that compared with cells transfected with empty vector, CDT1 overexpression did not dramatically change the cell cycle features of 3T3 cells in which ΔDboxGeminin was not induced.

**Supplementary Figure 10:**
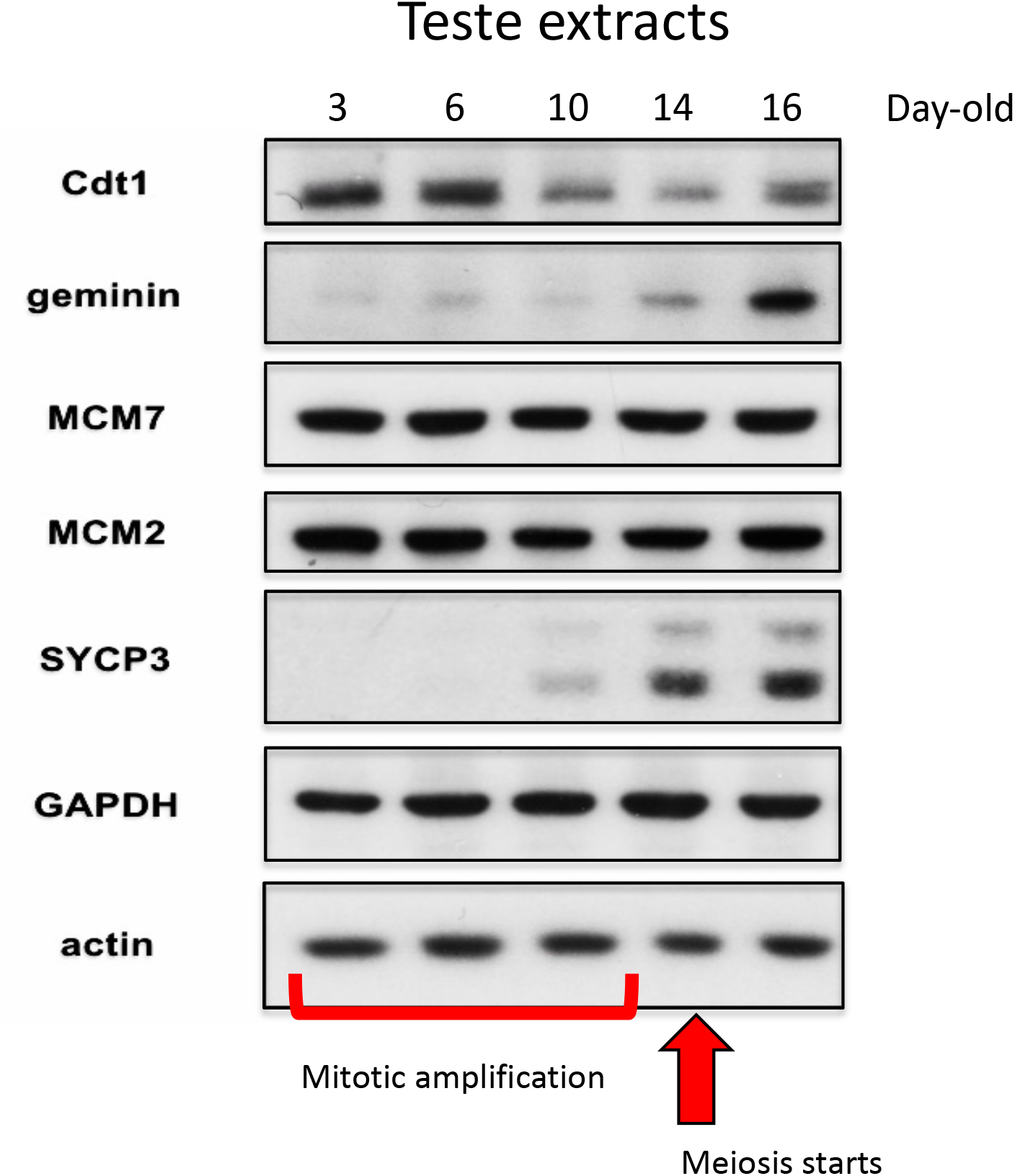
Geminin is upregulated during initiation of spermatogenesis in mice. Testes were dissected from Sv129/B6 mice at different days post-partum and the expression of the indicated proteins was assessed by western blot analysis. Actin and GAPDH were used as loading controls and SYCP3 to monitor meiosis onset.

**Supplementary Figure 11:**
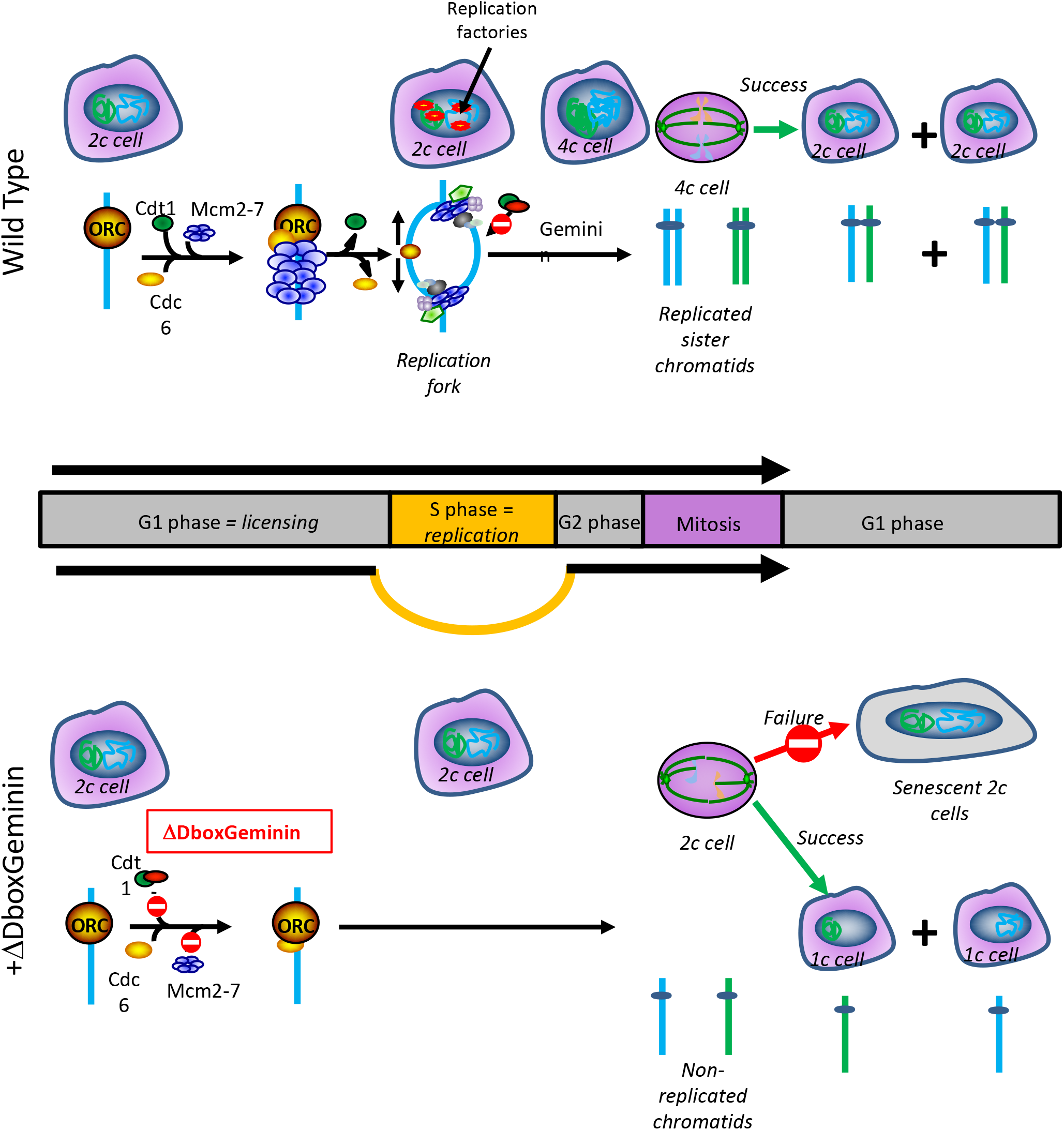
Summary of the different fates of normal somatic cells upon licensing inhibition. *Upper part:* Scheme of the temporal coordination between DNA replication and cell division in wild type diploid cells (2C DNA content). During the G1 phase of the cell cycle, the licensing reaction takes place, resulting in the binding of the DNA helicase MCM2-7 at DNA origins. This binding is dependent on the presence of the proteins ORC, CDC6 and CDT1. Upon S phase entry, replication origins are activated and DNA is replicated. Geminin starts to be synthesized and inhibits further binding of CDT1 to origins, preventing re-licensing and re-replication of DNA. When DNA replication is completed, cells enter into mitosis (4C DNA content), and the two replicated sister chromatids remain attached by the centromere until anaphase onset. *Lower panel*: Fates of non-transformed cells in which licensing is inhibited by overexpression of ΔDboxGeminin. The absence of the destruction box (Dbox) stabilizes geminin, which is normally degraded in G1 by APC/C. ΔDboxGeminin prevents binding of CDT1 to origins and inhibits pre-RC formation. Unlicensed cells enter mitosis with unreplicated chromatids (2C). The majority of these cells cannot complete mitosis, leading to the generation of one single cell with one (mitotic slippage) or two (cytokinesis failure) nuclei. These cells display nucleolar γH2AX staining and enter senescence. Nevertheless, about ¼ of unlicensed cells manage to divide and gives rise to two daughter cells, some of them with 1C DNA content.

## Notes

### Competing Interest Statement

The authors have declared no competing interest.

